# Data-Independent Acquisition and Quantification of Extracellular Matrix from Human Lung in Chronic Inflammation-Associated Carcinomas

**DOI:** 10.1101/2022.08.05.503012

**Authors:** Joanna Bons, Deng Pan, Samah Shah, Rosemary Bai, Chira Chen-Tanyolac, Xianhong Wang, Daffolyn R. Fels Elliott, Anatoly Urisman, Amy O’Broin, Nathan Basisty, Jacob Rose, Veena Sangwan, Sophie Camilleri-Broët, James Tankel, Philippe Gascard, Lorenzo Ferri, Thea D. Tlsty, Birgit Schilling

## Abstract

Early events associated with chronic inflammation and cancer involve significant remodeling of the extracellular matrix (ECM), which greatly affects its composition and functional properties. Using lung squamous cell carcinoma (LSCC), a chronic inflammation-associated cancer (CIAC), we optimized a robust proteomic pipeline to discover potential biomarker signatures and protein changes specifically in the stroma. We combined ECM enrichment from fresh human tissues, data-independent acquisition strategies, and stringent statistical processing to analyze ‘Tumor’ and matched adjacent histologically normal (‘Matched Normal’) tissues from patients with LSCC. Overall, 1,802 protein groups were quantified with at least two unique peptides, and 56% of those proteins were annotated as ‘extracellular’. Confirming dramatic ECM remodeling during CIAC progression, 529 proteins were significantly altered in the ‘Tumor’ compared to ‘Matched Normal’ tissues. The signature was typified by a coordinated loss of basement membrane proteins and small leucine-rich proteins. The dramatic increase in the stromal levels of SERPINH1/heat shock protein 47, that was discovered using our ECM proteomic pipeline, was validated by immunohistochemistry (IHC) of ‘Tumor’ and ‘Matched Normal’ tissues, obtained from an independent cohort of LSCC patients. This integrated workflow provided novel insights into ECM remodeling during CIAC progression, and identified potential biomarker signatures and future therapeutic targets.

**Statement of significance of the study:** The extracellular matrix (ECM) is a complex scaffolding network composed of glycoproteins, proteoglycans and collagens, which binds soluble factors and, most importantly, significantly impacts cell fate and function. Alterations of ECM homeostasis create a microenvironment promoting tumor formation and progression, therefore deciphering molecular details of aberrant ECM remodeling is essential. Here, we present a multi-laboratory and refined proteomic workflow, featuring i) the prospective collection of tumor and matched histologically normal tissues from patients with lung squamous cell carcinoma, ii) the enrichment for ECM proteins, and iii) subsequent label-free data-independent acquisition (DIA)-based quantification. DIA is a powerful strategy to comprehensively profile and quantify all detectable precursor ions contained in the biological samples, with high quantification accuracy and reproducibility. When combined with very stringent statistical cutoffs, this unbiased strategy succeeded in capturing robust and highly confident proteins changes associated with cancer, despite biological variability between individuals. This label-free quantification workflow provided the flexibility required for ongoing prospective studies. Discussions with clinicians, surgeons, pathologists, and cancer biologists represent an opportunity to interrogate the DIA digitalized maps of the samples for newly formulated questions and hypotheses, thus gaining insights into the continuum of the disease and opening the path to novel ECM-targeted therapies.

## 1. Introduction

Chronic inflammation-associated cancers (CIACs) account for one in four cancers worldwide and are responsible for more than 2 million deaths annually [1, 2]. Chronic inflammation can be caused by diverse biological, chemical and physical factors [3]. Cigarette smoke is one significant risk factor, that can promote lung cancers [3–5], including lung squamous cell carcinoma (LSCC) [6]. LSCC, a subtype of non-small cell lung cancer (NSCLC), accounts for about a third of all lung cancers. LSCC arises in the epithelial cells lining the bronchi, and it progresses through squamous metaplasia and dysplasia [7]. Interestingly, at the site of chronic injury, inflammation can favor cell plasticity and lead to a remodeling of the tissue microenvironment by altering stromal and extracellular matrix (ECM) homeostasis, which in turn can promote a malignant fate through poorly understood molecular mechanisms [1]. Additionally, the dynamic ECM remodeling can subsequently alter not only ECM composition and stiffness, but also initiate a cascade of biochemical and biophysical cues that affect, in turn, cell signaling. Ultimately, the ECM plays a key role in promoting tumor proliferation, invasion and metastasis [8–10], representing a crucial and promising intervention target for therapies: Could ‘repair’ of the ECM be a therapeutic intervention?

To enable in-depth ECM proteome characterization, Naba *et al.* pioneered the integration of proteomic and bioinformatic datasets to generate a database of ECM and ECM-associated proteins, referred to as the matrisome [11, 12]. The core matrisome includes collagens, ECM glycoproteins and proteoglycans, whereas the matrisome-associated proteins are composed of ECM-affiliated proteins, ECM regulators and secreted factors. Very recently, McCabe *et al.* reported an extensive mouse ECM atlas based upon the characterization of the ECM profiles of 25 organs by liquid chromatography-tandem mass spectrometry (LC-MS/MS) [13].

Advances in MS-based proteomics enable great opportunities to investigate ECM proteome remodeling in cancers and to identify novel protein biomarkers and therapeutic targets. In recent studies, aimed at uncovering ECM remodeling in various human cancers, data-dependent acquisition (DDA) label-free quantification approaches were employed to investigate glioblastoma and medulloblastoma [14], as well as gastric antrum adenocarcinoma [15]. DDA-tandem mass tag (TMT)-based quantification workflows were applied to analyze ECM from human pancreatic ductal adenocarcinoma [16] and to investigate metastasis in various mouse models of triple-negative mammary carcinoma [17, 18]. Finally, Naba *et al.* used isobaric tags for relative and absolute quantitation (iTRAQ)-based DDA quantification to analyze pancreatic islet ECM from a mouse model of insulinoma [19]. However, the semi-stochastic sampling and selection of precursor ions for MS/MS in DDA mode, in which the most abundant ions are selected for fragmentation during any given scan cycle, often lead to missing values and reproducibility challenges. For isobaric stable isotope labeling strategies, such as iTRAQ [20] and TMT [21], the labeled samples are typically pooled before DDA-MS acquisitions, implying that all samples are preferably collected and processed simultaneously, which is challenging for actively ongoing human studies, such as with prospective cancer patient studies. Other challenges in DDA-based isobaric labeling workflows are pairwise comparison ratio compression and quantification accuracy; however, strategies have been developed to address these challenges [22–24].

Alternatively, label-free data-independent acquisition (DIA) strategies performed on high-resolution, accurate-mass instruments represent a powerful tool to quantify ECM proteins across disease stages in prospective clinical cohorts, such as the one studied here. DIA relies on the systematic acquisition of MS/MS spectra for all detectable peptides contained in wide m/z isolation windows [25, 26]. Generated DIA MS/MS spectra are then interrogated using dedicated data processing strategies [27, 28], that typically rely on tissue-specific spectral libraries [29, 30], pan-species spectral libraries [31, 32] or library-free workflows, such as DIA-Umpire [33], DIA-NN [34] and directDIA embedded in Spectronaut software (Biognosys). In recent years, considerable efforts have been made to improve software algorithms [34–36], allowing to mine DIA data in-depth. DIA provides comprehensive and deep profiling of the proteome with highly reproducible and accurate quantification performances [37–39]. Numerous cancer and pre-clinical studies have been performed using DIA approaches [40, 41]. Krasny *et al.* reported the first application of the DIA/SWATH methodology to profile mouse liver and mouse lung matrisomes, and benchmarked the performances of DIA/SWATH vs. DDA [42]. The authors reported that DIA/SWATH achieved 54% more matrisomal protein identification and improved reproducibility performances compared to DDA-based analysis.

In this study, we present an efficient and robust multi-site workflow combining prospective collection of fresh human LSCC ‘Tumor’ and matched adjacent histologically normal (‘Matched Normal’) tissue specimens from 10 cancer patients, ECM enrichment at UCSF, label-free comprehensive DIA quantification and stringent statistical processing at the Buck Institute, and finally candidate verification by immunofluorescence-based immunohistochemistry (IHC) at UCSF (**Figure 1**). This workflow was applied to decipher ECM proteome remodeling in LSCC, thus allowing us to gain deeper mechanistic insights into how altered ECM can promote tumorigenesis, and to identify potential ECM targets whose modulation may restore ECM homeostasis and a microenvironment less permissive for malignancy.

**FIGURE 1.**
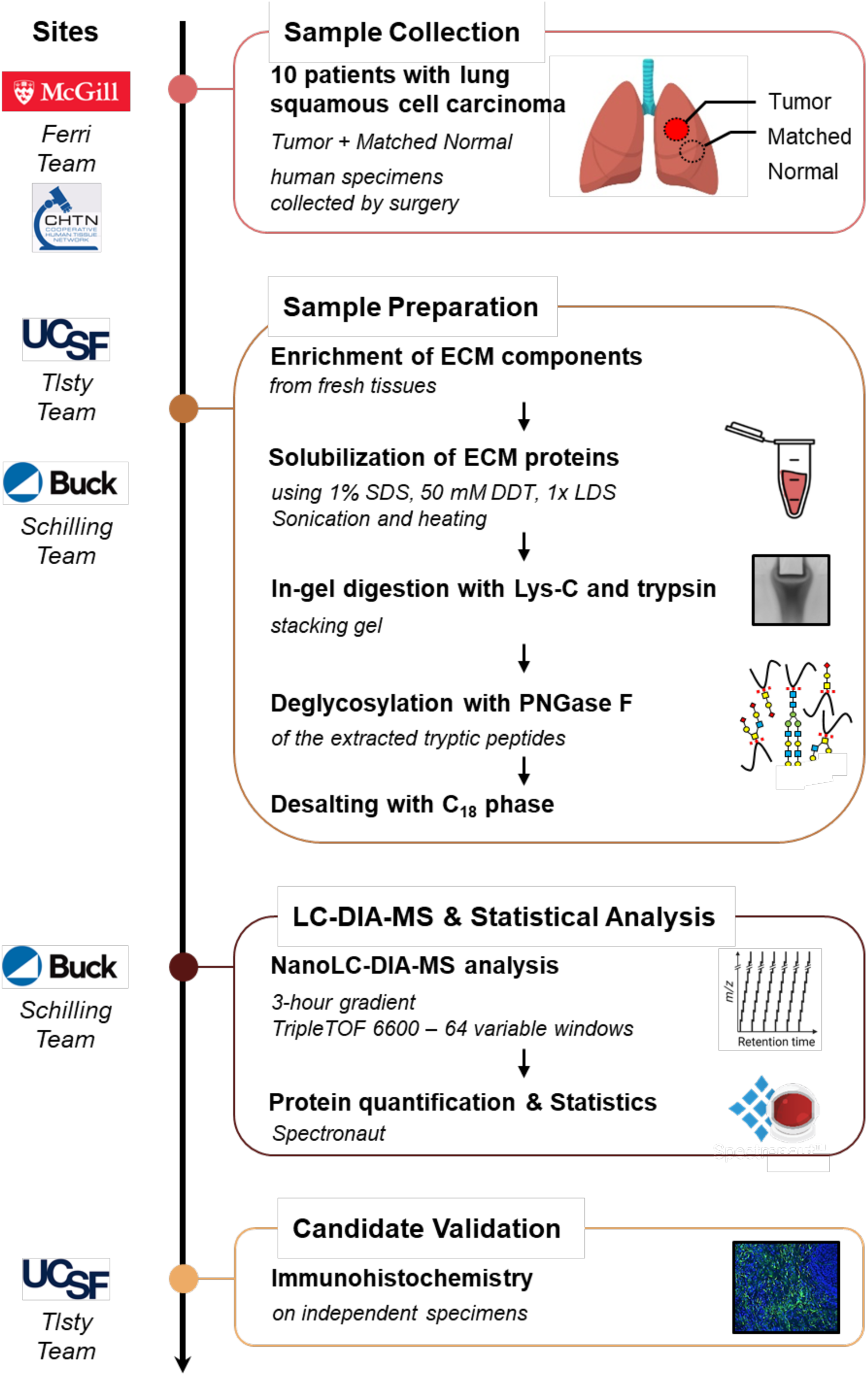
Study Design and Proteomic Pipeline for ECM Analysis of Lung Squamous Cell Carcinoma. ‘Tumor’ and ‘Matched Normal’ fresh tissue specimens were collected from 10 patients with lung squamous cell carcinoma by the Ferri team at McGill University Health Centre (QC, Canada) or through the Cooperative Human Tissue Network Western Division (TN, USA). Fresh samples in UW Cold Storage Solution were sent to the Tlsty team at UCSF (CA, USA) for enrichment for insoluble extracellular matrix (ECM) proteins. ECM-enriched samples were further processed by the Schilling team at the Buck Institute (Novato, CA); proteins were solubilized, in-gel digested with Lys-C and trypsin, and extracted proteolytic peptides were de-glycosylated with PNGase F. All resulting samples were analyzed in duplicate on a nanoLC-TripleTOF 6600 system (QqTOF) operated in data-independent acquisition (DIA) mode, and data were processed with Spectronaut (Biognosys). Finally, candidates were validated on independent cohorts by immunofluorescence-based immunohistochemistry by the Tlsty team.

## 2. Materials and Methods

### 2.1. Chemicals

LC-MS-grade acetonitrile (ACN) and water were obtained from Burdick & Jackson (Muskegon, MI). Reagents for protein chemistry, including sodium dodecyl sulfate (SDS), ammonium bicarbonate, iodoacetamide (IAA), dithiothreitol (DTT), sequencing-grade endoproteinase Lys-C, and formic acid (FA) were purchased from Sigma-Aldrich (St. Louis, MO). Sequencing-grade trypsin was purchased from Promega (Madison, WI). Glycerol-free PNGase F was purchased from New England BioLabs (Ipswich, MA).

### 2.2. Collection of Human Samples

#### Surgery Samples for ECM Analysis

Fresh tissue specimens corresponding to tumor (‘Tumor’) and histologically normal tissue adjacent to tumor (‘Matched Normal’) were collected from 5 female and 5 male consented patients diagnosed with lung squamous cell carcinoma at McGill University Health Centre (MUHC; Montreal, Quebec, Canada, 9 patients) or through the Western Division of the Cooperative Human Tissue Network (CHTN; Nashville, TN, USA, 1 patient). Sample collection was under protocols of the REB-approved biobank (study# 2007-856, lead investigator: Dr. Ferri) and used in the MUHC REB-approved CRUK STORMing project (local study# 2019-5039, lead investigator: Dr. Ferri), or under institutionally approved human subject protocol 10-01532 (University of California, San Francisco; UCSF). Tissue specimens were shipped overnight in transport medium (Belzer UW Cold Storage Solution; Bridge to Life Ltd., Northbrook, IL) to UCSF where they were processed for ECM proteomic analysis as described below. Information about each tissue specimen and patient, referred to as L01, L02, L03 …, and L10, is provided in **Table S1**.

#### Samples for Immunohistochemistry Validation

Formalin-fixed paraffin-embedded (FFPE) ‘Tumor’ and ‘Matched Normal’ lung specimens used for immunohistochemistry were provided by CHTN or by the Department of Pathology at UCSF under human subject protocol 10-01532 (UCSF). Information about each tissue specimen is provided in **Table S1**.

### 2.3. Enrichment for ECM Components

Fresh lung tissues were minced into small pieces, weighed and flash frozen for storage at -80 □C. The ECM fraction was isolated from the frozen tissues using the Compartmental Extraction Kit (Millipore, #2145) as per manufacturer’s protocol. Briefly, the tissues were homogenized in cold proprietary ‘buffer C’ using a bead mill homogenizer. ECM was extracted through step-wise washes with salt solutions and detergents to remove soluble proteins. About 1/10 of purified ECM was used to determine the purity and efficiency of the ECM protein enrichment by Western blot analysis. The ECM and other fractions were assessed using primary antibodies, specific for distinct cell fractions: anti-collagen I for ECM (Abcam, #ab138492), anti-beta 1 integrin for plasma membrane (Abcam, #ab179471), anti-hnRNPH1 for nucleus (Invitrogen, #27610), anti-GAPDH for cytosol (Abcam, #ab128915) and an anti-actin antibody for cytoskeleton (Sigma, #A5441) (**Figure S1**). The remaining ECM preparation was stored at -80□C for further quantitative proteomic analysis.

### 2.4. Solubilization of ECM Proteins

The extracted ECM pellets were solubilized by agitation for 10 minutes in a solution containing 1% SDS, 50 mM DTT and 1X NuPAGE lithium dodecyl sulfate (LDS) sample buffer (Life Technologies, Carlsbad, CA), followed by sonication for 10 minutes, and finally heating at 85 □C for 1 hour with agitation.

### 2.5. Protein Digestion and Desalting

Solubilized samples were run in pre-cast NuPAGE 4-12% gradient acrylamide Bis-Tris protein gels (Invitrogen) for 20 minutes to concentrate the proteins in a single band in the stacking gel. The gels were fixed with 50% methanol, 7% glacial acetic acid in water for 15 min and stained with GelCode Blue Stain (Thermo Fisher Scientific). For in-gel digestion, the gel bands were diced, collected in tubes, and dehydrated with a dehydration buffer (25 mM ammonium bicarbonate in 50% acetonitrile (ACN) and water). The gel samples were dried in a vacuum concentrator, reduced with 10 mM dithiothreitol (DTT) in 25 mM ammonium bicarbonate (pH 7-8) and incubated for 1 hour at 56 □C with agitation, then alkylated with 55 mM iodoacetamide (IAA) in 25 mM ammonium bicarbonate (pH 7-8), and incubated for 45 minutes at room temperature in the dark. The diced gel pieces were washed with 25 mM ammonium bicarbonate in water (pH 7-8), and then dehydrated again with the dehydration buffer. Samples were dried in a vacuum concentrator, after which the proteins were incubated with 250 ng of sequencing-grade endoproteinase Lys-C in 25 mM ammonium bicarbonate (pH 7-8) at 37 □C for 2 hours with agitation, followed by an overnight incubation with 250 ng sequencing-grade trypsin in 25 mM ammonium bicarbonate (pH 7-8) at 37 □C with agitation. Subsequently, the digested peptides were further extracted, as gel pieces were subjected to water and then two separate additions of a solution containing 50% ACN, 5% formic acid (FA) in water. After each addition of solution, the sample was mixed for 10 minutes, and then the aqueous digests from each sample were transferred into a new tube. These pooled peptide extractions were dried in a vacuum concentrator for three hours until completely dry. Proteolytic peptides were re-suspended in 100 µL of 25 mM ammonium bicarbonate in water (pH 7-8), and spot-checked to ensure a pH of 7-8. Subsequently, 3 µL (1,500 U) of glycerol-free PNGase F were added, and samples were incubated for 3 hours at 37 □C with agitation. This reaction was quenched with 10% FA in water for a final concentration of 1%, and spot-checked again to ensure a pH of 2-3. The quenched peptide samples were desalted using stage-tips made in-house containing a C_18_ disk, concentrated in a vacuum concentrator and re-suspended in aqueous 0.2% FA containing indexed retention time peptide standards (iRT, Biognosys, Schlieren, Switzerland) [43].

### 2.6. Mass Spectrometric Analysis

LC-MS/MS analyses were performed on an Eksigent Ultra Plus nano-LC 2D HPLC system (Dublin, CA) combined with a cHiPLC system directly connected to an orthogonal quadrupole time-of-flight (Q-TOF) SCIEX TripleTOF 6600 mass spectrometer (SCIEX, Redwood City, CA). The solvent system consisted of 2% ACN, 0.1% FA in H_2_O (solvent A) and 98% ACN, 0.1% FA in H_2_O (solvent B). Proteolytic peptides were loaded onto a C_18_ pre-column chip (200 μm × 6 mm ChromXP C18-CL chip, 3 μm, 300 Å; SCIEX) and washed at 2 μL/minute for 10 minutes with the loading solvent (H_2_O/0.1% FA) for desalting. Peptides were transferred to the 75 μm × 15 cm ChromXP C_18_-CL chip, 3 μm, 300 Å (SCIEX) and eluted at 300 nL/min with the following gradient of solvent B: 5% for 5 min, linear from 5% to 8% in 15 min, linear from 8% to 35% in 97 min, and up to 80% in 20 min, with a total gradient length of 180 min.

All samples were analyzed in technical duplicates by data-independent acquisition (DIA), specifically using variable window DIA acquisitions [25, 37, 38]. In these DIA acquisitions, 64 windows of variable width (5.9 to 90.9 m/z) are passed in incremental steps over the full mass range (m/z 400–1,250), as determined using the SWATH Variable Window Assay Calculator from SCIEX (**Table S2**). The total cycle time of 3.2 seconds includes a MS1 precursor ion scan (250 msec accumulation time), followed by 64 variable window DIA MS/MS segments (45 msec accumulation time for each). MS2 spectra were collected in “high-sensitivity” mode. The collision energy (CE) for each segment was based on the z=2+ precursor ion centered within the window with a CE spread of 10 or 15 eV.

### 2.7. DIA Data Processing with Spectronaut

All DIA data was processed in Spectronaut version 14.10.201222.47784 (Biognosys) using a pan-human library that provides quantitative DIA assays for 10,316 human proteins [31]. Data extraction parameters were selected as dynamic, and non-linear iRT calibration with precision iRT was selected. Identification was performed using a 1% precursor and protein q-value, and iRT profiling was selected. Quantification was based on the MS/MS peak area of the 3-6 best fragment ions per precursor ion, peptide abundances were obtained by summing precursor abundances and protein abundances by summing peptide abundances. Interference correction was selected and local normalization was applied. Differential protein abundance analysis was performed using paired t-test, and p-values were corrected for multiple testing, specifically applying group-wise testing corrections using the Storey method [44]. For differential analysis, very stringent criteria were applied: protein groups with at least two unique peptides, q-value ≤ 0.001, and absolute Log2(fold-change) ≥ 0.58 were considered to be significantly altered (**Table S3**).

### 2.8. Bioinformatic Analysis

The Pearson coefficients of correlation were determined between the different replicates using the cor() function of the stats package in R (version 4.0.2; RStudio, version 1.3.1093) and the abundances of all 1,802 quantifiable protein groups as input. Violin plots were generated using the ggplot2 package [45]. Partial least square-discriminant analysis (PLS-DA) of the proteomics data was performed using the package mixOmics [46] in R. An over-representation analysis was performed using ConsensusPathDB-human (Release 35, 05.06.2021) [47, 48] to determine which gene ontology (GO) terms were significantly enriched. The significantly 327 up- and 202 down-regulated protein groups were used as inputs and all quantified 1,802 protein groups were used as customized background proteome (**Table S3**). GO terms identified from the over-representation analysis were subjected to the following filters: q-value < 0.001 and term level ≥ 4. Dot plots were generated using the ggplot2 package [45] in R.

### 2.9. Immunohistochemistry and Quantification

FFPE tissue sections (5 μm thick) for six cases of LSCC and matched histologically normal lung tissues from two of these cases and four additional matched histologically normal lung tissues were deparaffinized and rehydrated in ethanol and water. Endogenous peroxidase was inactivated with 3% hydrogen peroxide for 10 min at room temperature. After antigen retrieval with citric acid buffer (10 min, 95°C), sections were blocked with background sniper (Biocare, BS966) then incubated for 1h at room temperature with a knock-out validated recombinant rabbit monoclonal anti-SERPINH1 (Hsp47) antibody (Abcam, #ab109117) diluted at 1/50 or 1/150 followed by an incubation for 1h at room temperature with a secondary goat Alexa Fluor^TM^ 488 antibody (ThermoFisher Scientific, #A11029) diluted at 1/1000. Tissue sections were counterstained with DAPI, mounted with VECTASHIELD HardSet Antifade Mounting Medium (Vector Laboratories, H-1400) and coverslipped. Sections were then imaged, specifically acquiring 5 images per each specimen, at a 20x magnification using a Keyence BZ-X800 series microscope. Images were processed with the ImageJ software, and Hsp47 abundance was quantified based on fluorescein isothiocyanate (FITC) staining using the QuPath (version 0.3.2) software. Plots and statistical analysis (Welch’s test) were processed with the Prism (version 9.3.1) software.

## 3. Results and Discussion

### 3.1. Efficient Proteomic Workflow for Human Lung Extracellular Matrix

To decipher changes that occur in the ECM of human chronic inflammation-associated lung squamous cell carcinoma, a refined multi-site experimental workflow was implemented, combining i) fresh human tissue collection immediately after resection and pathology assessment at McGill University (or CHTN Western Division in Nashville), ii) ECM isolation at UCSF, iii) proteomic analysis, including MS acquisition and statistical processing at the Buck Institute, and finally iv) biomarker candidate validation via immunofluorescence-based immunohistochemistry of an independent cohort of patients at USCF (**Figure 1**).

Approximately 50 mg of fresh lung tissue specimens from 10 patients with LSCC were collected by surgical resection, both at the tumor site and at a histologically normal site adjacent to the tumor, hereafter referred to as ‘Tumor’ and ‘Matched Normal’, respectively. Tissue annotations initially assigned by a surgeon were subsequently confirmed by a pathologist. The clinical traits of the 5 female and 5 male cancer patients with ages ranging from 57 to 78 years are displayed in **Table S1**. Fresh and never-frozen tissue specimens stored in cold UW solution were sent to USCF for further ECM enrichment by sequential fractionation based on solubility. The quality of the ECM protein enrichment was assessed using Western blotting assays by examining the abundance of representative proteins for specific cellular compartments/fractions: collagen I for the ECM fraction, β1 integrin for the membrane fraction, heterogeneous nuclear ribonucleoprotein H1 (hnRNP H1) for the nucleus fraction, glyceraldehyde-3-phosphate dehydrogenase (GAPDH) for the cytosol fraction and actin for the cytoskeleton fraction (**Figure S1**). Highly insoluble ECM was enriched as confirmed by the very abundant presence of collagen I observed in the anticipated ECM fractions. Representative markers for other cellular compartments were only minimally or not at all detected in the ECM fractions, documenting a high efficiency of the ECM isolation protocol and a high purity of the isolated ECM proteins. Isolated ECM from ∼50 mg of each original lung specimen, namely 10 ‘Tumor’ specimens and 10 ‘Matched Normal’ specimens, was sent for proteomic analysis to the Buck Institute. Insoluble ECM-enriched proteins were solubilized applying a rigorous procedure including a solution of anionic detergents (1% SDS, 0.5% LDS) and reducing agent (DTT), and samples were subjected to sonication (for 10 min) and extended heat treatment (85°C for 1 hour) to improve protein solubilization. After concentration in a short migration stacking SDS-PAGE gel, proteins were in-gel digested via sequential incubations with Lys-C and subsequently trypsin, ensuring a high proteolytic digestion efficiency. Extracted tryptic peptides were de-glycosylated using PNGase F to release N-linked glycans from the ECM proteins in order to facilitate the overall mass spectrometric analysis (**Figure 1**).

To circumvent mass spectrometric under-sampling as often observed in DDA-approaches, an efficient and comprehensive DIA-MS strategy was chosen. Briefly, DIA enables us to profile all detectable peptides contained in the samples through the unbiased acquisitions of multiplexed MS/MS spectra, thus providing a digitalized map of each of the 10 ‘Tumor’ and 10 ‘Matched Normal’ control samples. Duplicate injections were performed for each sample on a TripleTOF 6600 mass spectrometer operated in DIA mode, using a 64 variable window isolation scheme (**Table S2**). More specifically, each scan cycle was composed of one full range MS scan (m/z 400-1,250) and 64 MS/MS scans with isolation windows ranging between 5.9 m/z and 90.9 m/z, with smaller windows in highly populated m/z regions and wider windows in less populated m/z regions [25, 37, 38]. As a result, DIA MS/MS spectrum complexity is reduced and analyte specificity is increased. Collected DIA data were analyzed using a pan-human spectral library [31]. Although this publicly available pan-human library was originally generated from acquisitions on different LC-MS/MS systems (however, also SCIEX Q-TOF/TripleTOF systems), non-linear retention time calibrations using iRT regressions were achieved very efficiently (**Figure S2A**). Peptide quantification was performed by extracting fragment ion chromatograms from the DIA MS/MS spectra. Here, ∼11 data points per chromatographic peak were used on average, providing high quantification accuracy. This DIA-MS workflow resulted in the identification and quantification of 1,802 protein groups (**Figure 2A**) with at least two unique peptides at 1% false discovery rate (FDR) (**Table S3A**). The median protein abundance (based on peak area) span 4.95 orders of magnitude over the entire dataset (**Figure S2B**).

**FIGURE 2.**
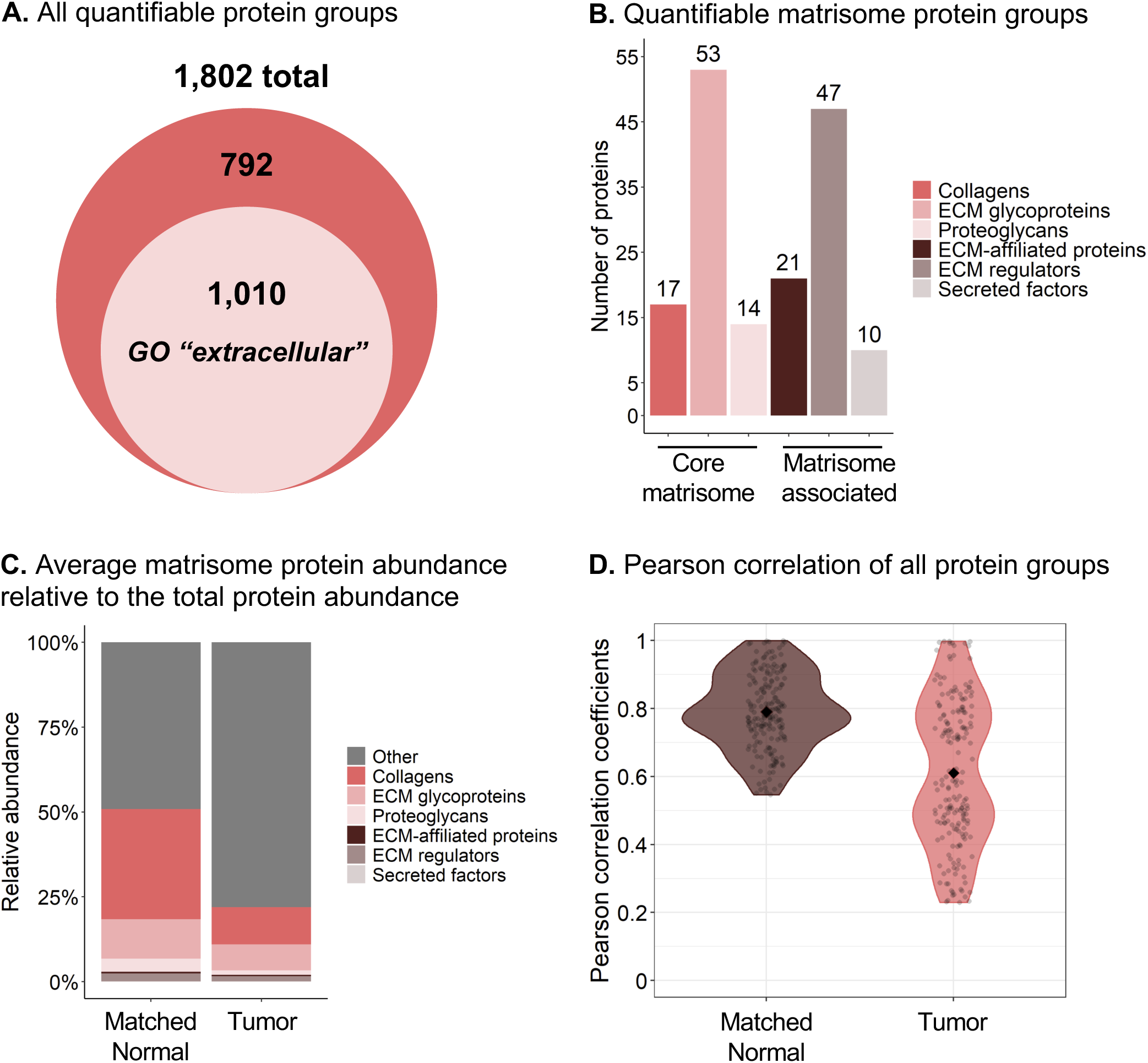
Proteomic Analysis of ECM and Matrisome Components from Human Lung Squamous Cell Carcinoma. (A) 1,802 protein groups with at least two unique peptides were identified, including 1,010 protein groups matching the Gene Ontology (GO) Cellular Component “extracellular” term. (B) 162 of these quantified protein groups are reported in the human MatrisomeDB [50]. (C) Average protein abundance relative to the total protein abundance of the matrisomal (colored) and non-matrisomal (grey) protein groups. Abundance is based on the MS/MS peak area of the 3-6 best fragment ions per precursor ion. Protein abundances were obtained with summing peptide/precursor abundances as described in the Methods section. (D) Violin plots of the Pearson coefficients of correlation between the ‘Matched Normal’ or ‘Tumor’ replicates. The Pearson correlation compares all MS acquisitions within one condition to each other (one by one). The filled diamonds represent the average value of the coefficients: 0.76 for the ‘Matched Normal’ group and 0.61 for the ‘Tumor’ group. High heterogeneity of the ‘Tumor’ ECM enrichments across cancer patients (right plot) contrasts with a more homogeneous profile for ‘Matched Normal’ ECM enrichments (left plot).

One powerful and unique aspect of this overall project is the prospective recruitment of patients, resulting in a continuous tissue collection, and subsequent proteomic analysis of tissue specimens. Using a label-free DIA-MS strategy represents a high advantage as it offers the flexibility required to prepare and acquire the tissue samples in an independent fashion and without any required sample pooling (as necessary for isobaric labeling strategies). To account for the technical variability, an efficient normalization method, based on a RT-dependent local regression model [49], was applied (**Figure S2C-D**). Briefly, assuming that the systematic bias is not linearly related to peptide abundances, the locally weighted scatterplot smoothing (LOWESS) algorithm is applied to perform a linear least squares regression on localized subsets of peptides (this algorithm is implemented into Spectronaut).

Of the 1,802 quantified protein groups obtained by analyzing the ECM-enriched fractions, 1,010 protein groups (56%) matched the gene ontology (GO) cellular component ‘extracellular’ (**Figure 2A**). Specifically, 162 protein groups are reported as ECM and ECM-associated proteins in the human MatrisomeDB database [50] (**Figure 2B**), with 17 collagens, 53 glycoproteins, 14 proteoglycans, 21 ECM-affiliated proteins, 47 ECM regulators and 10 secreted factors (**Table S3A**). Strikingly, although matrisomal proteins represented 9% of all quantified protein groups in the dataset, their peak area-based abundance accounted for 51% of the total protein abundance in the ‘Matched Normal’ group and 22% in the ‘Tumor’ group (**Figure 2C**). Indeed, over 50% of the quantified matrisomal protein groups (84/162) were present among the first most abundant protein groups quartile, with, for instance, collagen alpha-1(VI) chain, collagen alpha-2(VI), collagen alpha-3(VI) chain, fibronectin, and vitronectin in the top 10 most abundant protein groups (**Table S2B**). This highlights the efficient enrichment for ECM proteins achieved using this workflow. **Figure 2D** displays violin plots of the Pearson coefficients of correlation between the different replicates of each of the ‘Tumor’ and ‘Matched Normal’ groups. The increased variability in the ‘Tumor’ group compared to the ‘Matched Normal’ group revealed here that ECM enrichments from tumors are biologically highly heterogeneous across cancer patients, while the ECM from the matched histologically normal lung tissues appeared much more homogeneous across individuals.

Overall, these DIA-MS results already demonstrated the high ECM enrichment efficiency and quantification capabilities achieved by this presented workflow to comprehensively quantify ECM proteins, enriched from fresh clinical human tissue specimens. This label-free DIA strategy provides comprehensive, reproducible, sensitive, and accurate quantification of the ECM-enriched samples, while being compatible with the prospective and continuous addition of new samples to the cohort.

### 3.2. Human Lung Squamous Cell Carcinoma Features ECM Remodeling

By investigating the quantitative DIA-MS results more closely, it was obvious and interesting to discover that both ‘Tumor’ and ‘Matched Normal’ groups were quite distinct, and could be clearly clustered apart using a supervised clustering analysis by partial least squares-discriminant analysis (PLS-DA) (**Figure 3A**). Notably, distinct clustering was observed for specimens collected from both male and female patients (**Figure S3**). To explore the remodeling of ECM associated with LSCC, very stringent significance thresholds, specifically with q-value ≤ 0.001 and absolute Log_2_(fold-change) ≥ 0.58, were applied. The differential analysis of all 1,802 protein groups resulted in 529 significantly changing proteins comparing ‘Tumor’ to ‘Matched Normal’ samples. Specifically, this analysis revealed 327 significantly up-regulated protein groups and 202 significantly down-regulated protein groups (in ‘Tumor’ vs. ‘Matched Normal’) as shown in **Figure 3B** and **Table S3B**. Among the significantly changing proteins, 49 protein groups are well-known components of the core matrisome: 12 collagens, 29 ECM glycoproteins, and 8 proteoglycans, whereas 17 protein groups are matrisome-associated proteins: 4 ECM-affiliated proteins, 10 ECM regulators, and 3 secreted factors [50] (**Figure 3C**; **Figure S4**; **Table S3B**).

**FIGURE 3.**
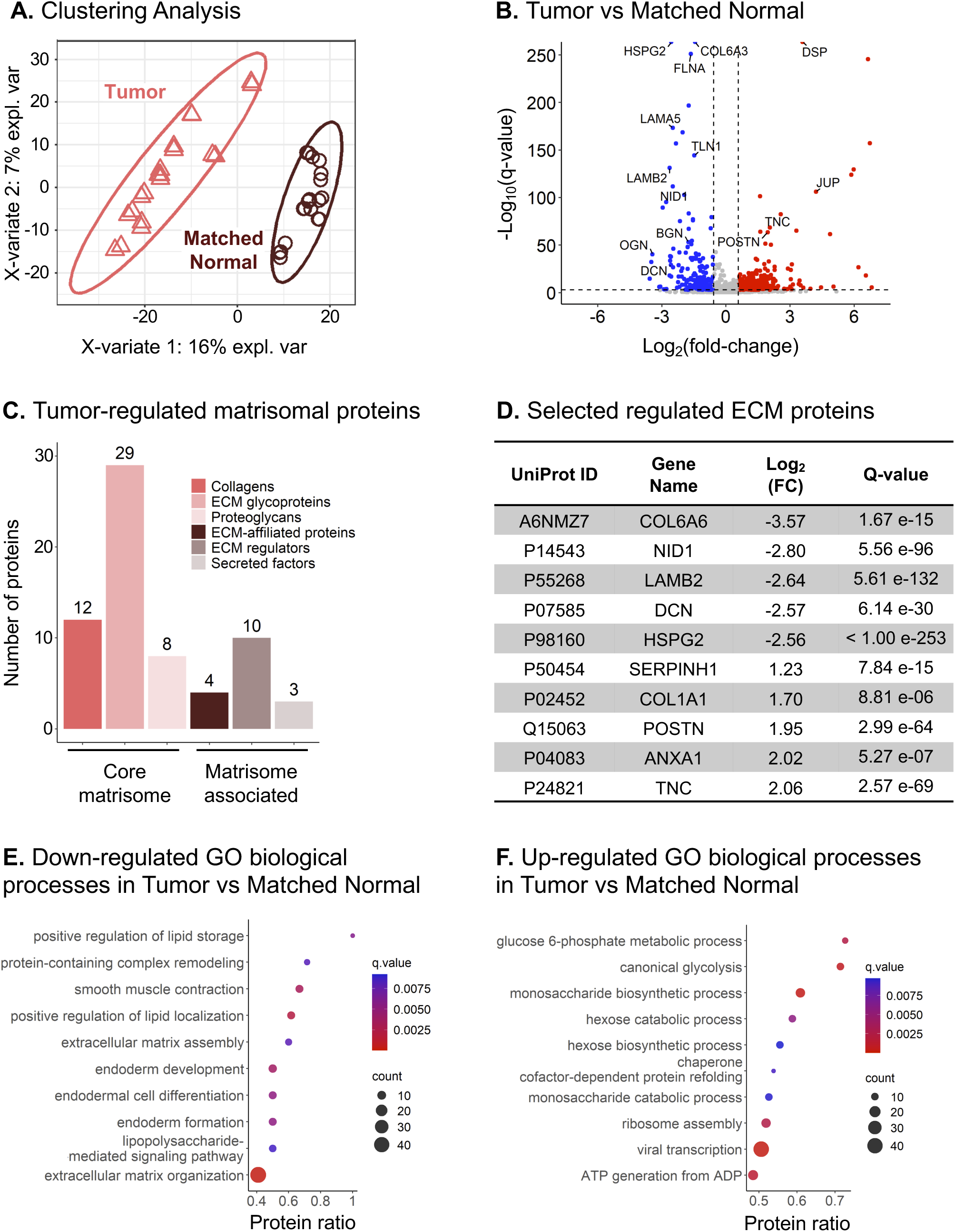
‘Complete’ Remodeling of the ECM in Lung Squamous Cell Cancer. (A) Supervised clustering analysis using partial least squares-discriminant analysis (PLS-DA) performed on protein groups quantified in the ‘Matched Normal’ (brown) and ‘Tumor’ (pink) samples. (B) Volcano plot of the 1,802 quantified protein groups showing 202 down-regulated and 327 up-regulated protein groups for ‘Tumor’ vs. ‘Matched Normal’ comparison. (C) 66 significantly altered protein groups are reported in the human MatrisomeDB [50] and are listed in **Figure S4**. Specific proteins from these significantly altered protein groups are listed in (D). (E-F) Dot plots showing the ConsensusPathDB [47, 48] Gene Ontology (GO) biological processes enriched for protein groups significantly down-regulated (E) and up-regulated (F) in ‘Tumor’ vs. ‘Matched Normal’.

Interestingly, ECM proteins down-regulated in ‘Tumor’ vs. ‘Matched Normal’ included collagen alpha-6(VI) chain (COL6A6), nidogen-1 (NID1), laminin subunit β2 (LAMB2), decorin (DCN), and perlecan (HSPG2), which are components of the basement membrane, a thin and specialized ECM layer. In contrast, serpin family H member 1/heat shock protein 47 (SERPINH1/Hsp47), a member of the serine protease inhibitor (serpin) family, collagen alpha-1(I) chain (COL1A1), periostin (POSTN), annexin A1 (ANXA1), and tenascin-C (TNC) were significantly up-regulated (**Figure 3D**), when comparing ‘Tumor’ to ‘Matched Normal’.

Of these significantly up-regulated protein groups, several protein candidates could potentially be highly relevant in the context of cancer and disease progression. For example, tenascin-C, which showed a 4.16-fold-increase in ‘Tumor’ vs. ‘Matched Normal’ with q-value = 2.57e-69 (**Figure 3D**), is a glycoprotein and member of the tenascin family. Tenascin-C is barely expressed in adult tissues, except in specific niches, such as at inflammation sites and in the stroma of solid tumors, where it is highly abundant [51]. In NSCLC, tenascin-C may participate in tumor immune evasion, progression, and recurrence via a mechanism involving the inhibition of tumor-infiltrating lymphocyte proliferation and interferon-γ secretion [52].

Additionally, annexin A1, up-regulated by a factor 4.06 in ‘Tumor’ vs. ‘Matched Normal’ with q-value = 5.27e-7 (**Figure 3D**), is a member of the Ca^2+^-regulated phospholipid-binding protein superfamily, involved in various cellular processes, such as inflammation, proliferation regulation, apoptosis, and tumorigenesis [53]. Notably, Annexin A1 appears as prognostic factor for longer overall survival in LSCC by suppressing metastasis, but not cancer cell proliferation [54].

Tumor ECM is known to be mechanically stiffer and exhibit higher tension compared to the ECM of healthy tissues. Collagens, whose organization relies on sophisticated crosslinking networks, largely contribute to this phenomenon [8], suggesting that COL1A1, here up-regulated by a factor 3.25 in ‘Tumor’ (q-value = 8.81e-6) (**Figure 3D**), could participate in ECM stiffening. Moreover, COL1A1 is associated with hypoxia in NSCLC [55]. This protein was also reported to correlate with late LSCC progression, and it appears as a potential biomarker of metastasis to lymph nodes [56], poor prognosis and chemoresistance [57] in LSCC.

Periostin was also significantly up-regulated (3.86-fold) in ‘Tumor’ vs. ‘Matched Normal’ with q-value = 2.99e-64 (**Figure 3D**). Periostin is a key player in ECM structure and organization, particularly for collagen fibrillogenesis, and it interacts with other proteins, such as integrins, fibronectin and tenascin [58, 59]. This protein is primarily expressed by cancer-associated fibroblasts (CAFs), located in the stromal microenvironment, and has been implicated in LSCC progression, tumor cell proliferation and migration [58, 60]. Specifically, Ratajczak-Wielgomas *et al.* reported that periostin in LSCC cancer cells could modulate the expression of the proteinase MMP-2, which may further regulate tumor cell invasion, and that periostin expression correlates with the incidence of lymph node metastases [61]. Moreover, the authors showed a putative interaction between cancer cells and stromal CAFs, which could promote cell invasion. Finally, periostin is associated with poor prognosis and tumor grade [58, 60], and correlates with the incidence of lymph node metastasis [60, 61].

To determine the biological processes altered in LSCC, an over-representation analysis was performed with ConsensusPathDB database [47, 48]. The gene ontology (GO) analysis revealed that processes related to lipid storage and localization, lipopolysaccharide-mediated signaling pathway, protein-containing complex remodeling, ECM organization, and endoderm development were down-regulated (**Figure 3E**) in ‘Tumor’ vs. ‘Matched Normal’ comparison. In contrast, processes related to glucose-6-phosphate metabolism, glycolysis, ATP generation, chaperon cofactor-dependent protein folding, and ribosome assembly were up-regulated (**Figure 3F**). The alterations of these biological processes clearly revealed aberrant ECM remodeling as well as alterations of metabolism in tumor cells. More specifically, sugar metabolism is reprogrammed in cancer cells as characterized by an exacerbated glucose uptake and a strong increase in lactate production, a phenomenon known as ‘the Warburg Effect’ [62], which may further impact the tumor microenvironment by favoring cell invasion and immunotolerance [63]. Down-regulation of lipid-related processes could be linked to alterations in the plasma membrane organization and/or of the lipid metabolism [64]. Proteins associated with such biological processes include apolipoprotein A-I (APOA1), a component of high-density lipoproteins involved in the transport of cholesterol, required for tumor cell viability, which in turn might promote tumor progression [65]. Another example is caveolin-1 (CAV1), which can undergo autophagic degradation in CAFs to protect adjacent epithelial tumor cells against apoptosis.

### 3.3. Changes in Basement Membrane Proteins, Small Leucine-Rich Proteins, Serpins, Desmosomal Proteins and Keratins

To decipher how ECM is remodeled in LSCC, the dataset was investigated by focusing on proteins involved in ECM structure and organization, as well as on specific protein families. **Figure 4** and **Figure S5** display heatmaps of the ‘Tumor’ vs. ‘Matched Normal’ significant fold-changes for the ECM abundance of basement membrane proteins, small leucine-rich proteins (SLRPs), serpins, desmosomal proteins, and keratins measured for each patient. Interestingly, very robust and strong signatures were observed for each of the patients.

**FIGURE 4.**
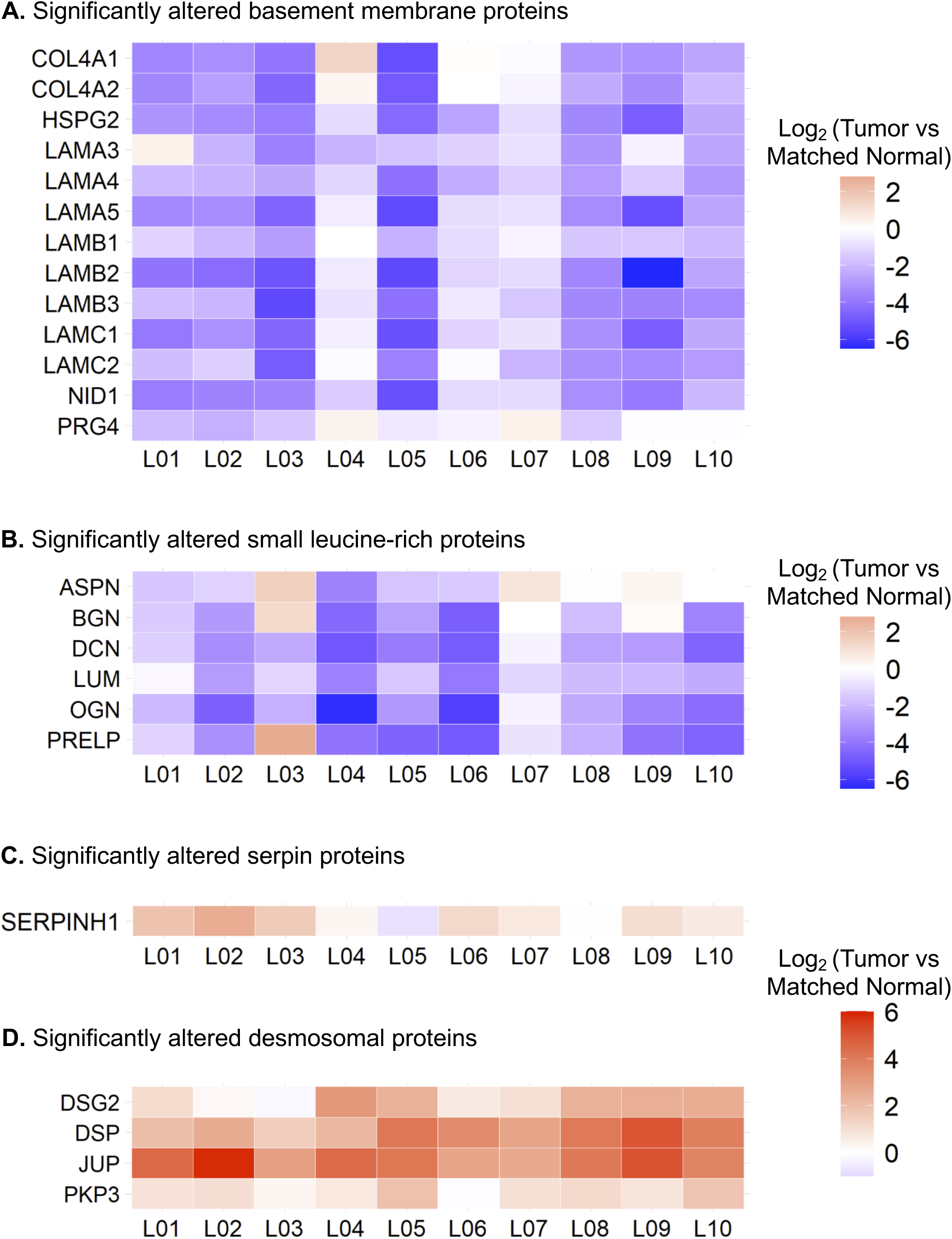
Alteration of Core Protein Signatures in Lung Squamous Cell Carcinoma across Patients. Heatmap showing log_2_-fold protein changes (‘Tumor’ vs. ’Matched Normal’) for each individual LSCC patient referred to as L01, L02, L03 …, and L10 assessing Core Protein Signatures. (A) Thirteen basement membrane proteins and (B) six small leucine-rich proteins were significantly down-regulated in ‘Tumor’ vs. ‘Matched Normal’, while (C) SERPINH1 (Hsp47) and (D) desmosomal proteins were significantly up-regulated across patients. For all displayed proteins, all Q-values (not displayed) were smaller than 9.38e-8, when comparing ‘Tumor’ to ‘Matched Normal’ group (**Table S3**).

Thus, abundance of basement membrane proteins, including collagen alpha-1(IV) chain (COL4A1), collagen alpha-2(IV) chain (COL4A2), collagen alpha-1(XVIII) chain (COL18A1), heparan sulfate proteoglycan 2/perlecan (HSPG2), eight laminin subunits – subunit α3 (LAMA3), α4 (LAMA4), α5 (LAMA5), β1 (LAMB1), β2 (LAMB2), β3 (LAMB3), γ1 (LAMC1), and γ2 (LAMC2), nidogen-1 (NID1), and proteoglycan 4 (PRG4), decreased in ‘Tumor’ compared ‘Matched Normal’ lung samples (**Figure 4**). Agrin (AGRN) and nidogen-2 (NID2) were also highly confidently down-regulated and close to the fold-change cutoff (q-value < 0.001 for both proteins; **Table S3**). The basement membrane is a thin layer of the ECM located at the base of polarized epithelial cells. It is composed of two independent networks of laminins and type IV collagens linked together by the glycoprotein nidogen and the heparan sulfate proteoglycan perlecan, as well as additional proteins including agrin and type XVIII collagens, which tether growth factors [66]. The basement membrane is involved in supporting the tissue architecture, maintaining cell polarity and signal transmission to epithelial cells via integrin receptors and segregating the epithelium from the stroma. Thus, the observed loss of basement membrane proteins is in accordance with its breaching, which is required for tumor cells to invade the stroma [67, 68].

In a similar fashion, a coordinated loss of SLRPs was observed in LSCC, specifically for asporin (ASPN), biglycan (BGN), decorin (DCN), lumican (LUM), mimecan (OGN), and prolargin (PRELP) (**Figure 4**). SLRPs represent a subgroup of proteoglycans, that is divided into four classes based on gene and protein homology: ASPN, BGN, and DCN belong to class I, LUM and PRELP to class II, and OGN to class III [69]. SLRPs are involved in various processes including ECM assembly regulation, collagen fibrillogenesis, sequestration of growth factors, cell-matrix interactions, and cell behaviors by interacting with plasma membrane receptors, such as toll-like receptors, tyrosine kinase receptors, and other matrisomal factors [70]. For example, decorin acts as a tumor suppressor by limiting tumor growth, angiogenesis, tumor cell mitophagy, and regulating the immune and inflammatory response [71]. While decorin and biglycan are the closest SLRPs, biglycan shows opposite activities by promoting inflammation, angiogenesis, tumor cell proliferation, migration, and metastasis, although tumor suppressive effects of this SLRP were also reported [71, 72]. Lumican binds to collagens to prevent degradation by proteinases, such as matrix metalloproteinases (MMPs), and is observed to have both pro- and anti-tumoral properties by regulating cell proliferation and invasion [72, 73]. Understanding the effects and interplay of these different SLRPs, that show significantly lower abundance in LSCC ‘Tumor’ vs. ‘Matched Normal’, may thus be of high relevance.

Strikingly, the overall loss of the core matrisome and matrisome-associated proteins in the ‘Tumor’ vs. ‘Matched Normal’ stroma, as illustrated for the basement membrane proteins and SLRPs, explains the >2-fold decrease of the matrisomal protein abundance relative to the total protein abundance in the ‘Tumor’ group compared to the ‘Matched Normal’ group (**Figure 2C**).

In our study, the observed alterations in protein abundance in the ECM for serine protease inhibitors (serpins) in LSCC were different for different family members: serpin family B member 6 (SERPINB6) was significantly down-regulated in ‘Tumor’ vs. ‘Matched Normal’ (ratio = 0.59 with q-value = 2.24e-4), whereas both serpin family B member 5/maspin (SERPINB5) and SERPINH1 (also referred to as heat shock protein 47, HSP47) were significantly up-regulated in ‘Tumor’ vs. ‘Matched Normal’ (SERPINB5: ratio = 1.91 with q-value = 8.20e-9; SERPINH1: ratio = 2.35 with q-value = 7.84e-15) (**Figure 4**; **Figure S5**). SERPINB6 interacts with cathepsin G in monocytes and granulocytes to inhibit this inflammation-related protein and interacts with other trypsin-like proteases as well [74]. SERPINB5 has a tumor suppressive activity [74]. In LSCC, this protein may be associated with cancer development by regulating the p53 signaling pathway [75]. SERPINH1 is an endoplasmic reticulum protein with chaperone activity which ensures proper folding and ultimately conformation of type I procollagen trimer [76]. SERPINH1 is dysregulated in a large number of cancers and might play a role in tumor immunity [77]. For instance, in breast cancer, Hsp47/SERPINH1 is a key player in cancer progression by promoting the secretion and deposition of ECM proteins, *e.g.* collagens and fibronectin [78], as well as of metastasis by regulating the cancer cell-platelet interaction via a collagen-dependent mechanism [79].

The ECM-enriched fractions contain residual highly insoluble remnants of stromal and epithelial cells, that were reproducibly co-isolated with ECM proteins. This was revealed by robust observations of keratins, which are epithelium intermediate filaments, involved in cell mechanical stability and integrity. Out of the 17 significantly altered keratins, 15 were up-regulated in ‘Tumor’ vs. ‘Matched Normal’ (**Figure S5**; **Table S3**), namely keratin, type I cytoskeletal 4, 5, 6A, 6B, 7, 8, 13, 14, 15, 16, 17, 18, 19, 75 and 80 (KRT4-8 and KRT13-19, KRT75 and KRT80). Two keratins, keratin, type I cytoskeletal 2 epidermal (KRT2) and keratin, type I cytoskeletal 9 (KRT9) were down-regulated. The observed global upregulation of keratins in this study highlights the strong keratinization process observed during LSCC, as also previously reported in laser microdissected tumor cells [80]. Interestingly, keratinization might be associated with smoking, a risk factor of lung CIAC, and keratinization correlates with poor clinical outcome in LSCC [81].

In addition, a coordinated, significant gain of desmosomal proteins was observed in ‘Tumor’ vs. ‘Matched Normal’, with <u>the up-regulation of desmoglein-2 (DSG2), junction plakoglobin (JUP), desmoplakin (DSP) and plakophilin-3 (PKP3)</u> (**Figure 4**; **Table S3**). Furthermore, plakophilin-1 and 2 (PKP1, PKP2) and desmocollin-2 (DSC2) were also up-regulated, however with slightly lower fold-change or just above the q-value cutoff of 0.001 (PKP1: ratio = 1.18 with q-value = 7.72e-9; PKP2: ratio = 1.56 with q-value = 0.0012; DSC2: ratio = 1.42 with q-value = 5.81e-4) (**Figure S5**). Desmosomes are intercellular junctions located on the lateral sides of plasma membranes and involved in cell-cell adhesion and resistance to mechanical stress. These structures are composed of desmosomal cadherins, such as desmocollins and desmogleins, desmosomal plaque proteins, such as desmoplakins and plakoglobins, as well as plakophilins. Notably, PKP1, KRT15 and DSG3 have been recently validated as novel markers to differentiate LSCC and lung adenocarcinoma, another NSCLC subtype, PKP1 and DSG3 being associated with poor prognosis [82]. In addition, PKP1 overexpression contributes to cell proliferation and survival in LSCC by positively regulating MYC translation [83]. Our study, presented here, highlights the relevance of the desmosomal protein assembly as part of LSCC.

Altogether, the significant and robust changes in the ECM of tumor tissues revealed a conserved signature in LSCC characterized by the concomitant loss of basement membrane proteins and SLRPs and the increase in SERPINH1, as well as other significant changes relative to keratin and desmosome protein family members.

### 3.4. SERPINH1 ECM Levels are Dramatically Increased in LSCC

To validate the highly significant up-regulation of SERPINH1 observed in LSCC ‘Tumor’ vs. ‘Matched Normal’ as determined by the mass spectrometric ECM analysis (2.35-fold increase; q-value of 7.84e-15), we employed an orthogonal method relying on IHC. Immunofluorescence-based IHC was conducted on an independent cohort of patients with LSCC from CHTN and UCSF. Samples for IHC were tumor tissues from six cancer patients with LSCC, two matched normal lung tissues from two of the cancer patient cases, and four additional histologically normal lung tissue specimens adjacent to lung cancers (**Table S1**). **Figure 5A** displays representative IHC images, demonstrating the strong up-regulation of SERPINH1 in cancer stroma in LSCC tissues, while it was barely detected in ‘Matched Normal’ tissues. Quantification was based on the percentage of area with positive FITC staining in five independent images for each specimen (**Figure 5B**). The quantitative analysis of stained tissue sections confirmed the dramatic up-regulation of SERPINH1 in LSCC ‘Tumor’ tissues compared to ‘Matched Normal’ tissues. A mean of 0.39% SERPINH1-positive area was obtained in the six ‘Matched Normal’ specimens; SERPINH1 increased 10.36-fold (p-value = 0.0116) to 4.05% SERPINH1-positive area in the six ‘Tumor’ specimens. While SERPINH1 was previously reported as a factor influencing tumor immunity and metastasis [77–79], further investigations need to be performed to determine the biological significance of SERPINH1 in LSCC and perhaps more broadly in multiple CIAC tumor types.

**FIGURE 5.**
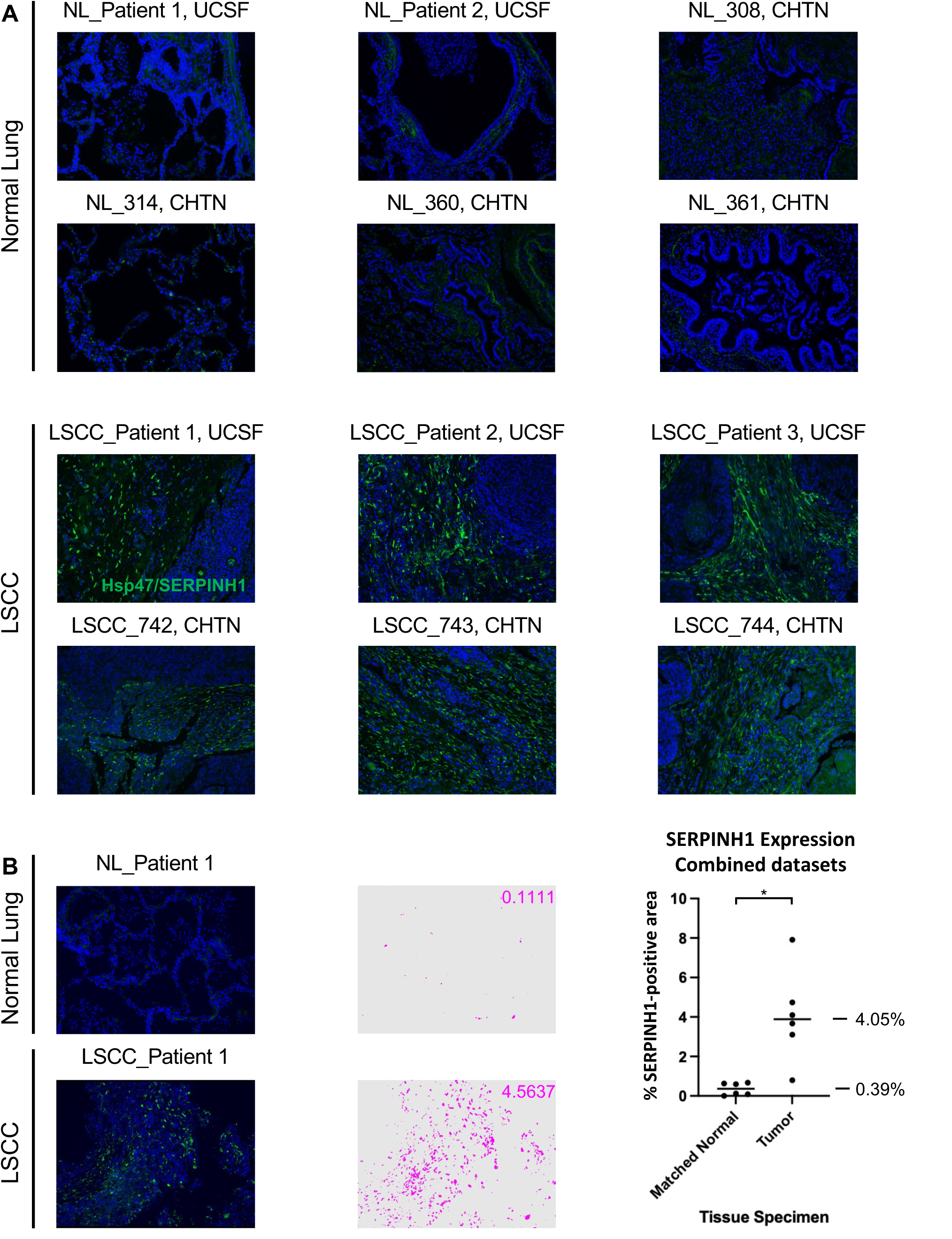
Dramatical Upregulation of SERPINH1 (Hsp47) in Lung Squamous Cell Carcinoma (LSCC) Compared to Histologically Normal Lung (NL) Tissues. (A) Six cases of lung squamous cell carcinomas and matched histologically normal lung tissues from two of these cases and four additional histologically normal lung tissue specimens adjacent to lung cancers were probed for SERPINH1 level by immunofluorescence-based immunohistochemistry (IHC). (B) Left: SERPINH1 level was quantified based on the percentage of area with positive (FITC; green) staining in 5 independent images per specimen (pixels with positive staining above baseline threshold/total number of pixels per image). An example of pseudo-colorized positive area (pink) is shown for a matched set of specimens. Right: Plot corresponding to averaged values of positive staining of 5 images for each of 12 human specimens (6 ‘Matched Normal’ and 6 ‘Tumor’). Statistical analysis was carried out as described in the Methods section. Magnification: 20x.

## 4. Concluding Remarks

In summary, our work presents an efficient, robust and multi-laboratory proteomic workflow to gain in-depth insights into ECM remodeling in LSCC by prospectively collecting fresh tumor and patient-matched histologically normal tissue adjacent to tumor from patients, enriching for insoluble ECM components, performing a refined ECM protein solubilization, applying comprehensive label-free DIA quantification and stringent statistical filtering. The unbiased DIA strategy offers the possibility to capture highly confident and robust protein changes, overcoming the biological individual-to-individual variability, while providing the flexibility required to prospective studies and sample collection scheduling. It is worth noting that one can consider applying this ECM proteomic workflow to any type of cancer or, more globally, diseases of interest. Although this study was conducted on cohorts with a limited number of patients, namely 10 patients for the discovery ECM proteomic analysis, it suggests that potential protein candidates can be further assessed and validated by orthogonal methods, such as Western blotting and IHC assays, on independent cohorts thus generalizing the observations. In this study, the application of immunofluorescence-based IHC confirmed in a second cohort the dramatic increase in Hsp47/SERPINH1 abundance identified by MS in the tumor stromal microenvironment of a first cohort of LSCC patients. In addition, the application of label-free global DIA strategy for the discovery step is an asset for the easy and efficient development of targeted parallel reaction monitoring (PRM) assays on similar MS platforms as validation step and further translation into true clinical cohort measurements. Moreover, as small amounts of material are needed for the presented workflow, MS-based proteomics can be further integrated with additional - omics technologies, such as (epi)genomics, transcriptomics, and CODEX, performed on the same tissue specimens in order to achieve a multifaceted tissue assessment. The combination of this compelling approach with regular discussions between surgeons, pathologists, cancer biologists and -omics scientists represents a cornerstone to formulate new hypotheses, and thus to gain deeper mechanistic insights into the continuum of disease processes and identify novel and promising stromal-targeted therapies.

## Supporting information

Figure S4 - page 2

## Acknowledgments

We acknowledge the support of instrumentation for the TripleTOF 6600 from the NIH shared instrumentation grant 1S10 OD016281 (Buck Institute). We thank Ms. Kerry Wiles and Mr. Erik Brooks at the Western Division of the Cooperative Human Tissue Network (Nashville, TN, USA) for providing some of the specimens used in this study. Dr. Birgit Schilling thanks Dr. Eric Verdin for his generous support. This work was generously supported by Cancer Research UK (CRUK) awards A27145 to Dr. Thea D. Tlsty (UCSF) and A29071 to Dr. Lorenzo Ferri (MUHC) as part of STORMing Cancer (Cancer Grand Challenge).

## Conflicts of Interest

The authors have declared no conflict of interest.

## Abbreviations

CAF: cancer-associated fibroblast
CE: collision energy
CIAC: chronic inflammation-associated cancer
DDA: data-dependent acquisition
DIA: data-independent acquisition
ECM: extracellular matrix
FA: formic acid
FFPE: formalin-fixed paraffin-embedded
IAA: iodoacetamide
IHC: immunohistochemistry
LDS: lithium dodecyl sulfate
LSCC: lung squamous cell carcinoma
NSCLC: non-small cell lung cancer
PLS-DA: partial least squares-discriminant analysis
RT: retention time
SLRP: small leucine-rich protein
TMT: tandem mass tag

## Supplementary Figure Legends

**FIGURE S1: ECM Isolation and Quality Control.** Approximately 50 mg of tissue from the ‘Matched Normal’ and ‘Tumor’ tissues of lung squamous cell carcinomas, obtained by the Ferri team (MUHC) or the CHTN Western Division, was homogenized and the different compartment fractions were extracted based on the instruction of a compartmental protein extraction kit (Millipore, #2145). For each sample, 12 µL of total tissue extract, 24 µL of intermediate fractions, and 1/10 of ECM isolated from 50 mg of tissue were used to examine levels of representative proteins for each compartment. Images of ‘Matched Normal’ (N) and ‘Tumor’ (T) tissues from MUHC patients L01 and L05 are shown. Insoluble ECM was enriched as documented by the high levels of Collagen I in the expected ECM fractions. Representative markers for other cellular compartments (cytoplasmic: GAPDH; nuclear: hnRNP H1; membrane: β1 integrin; cytoskeleton: Actin) were barely detected in these ECM fractions supporting high purity of the isolated ECM fraction. Isolated ECM from 50 mg of each tissue was processed for further proteomic analysis.

**FIGURE S2: Quality Control and Performance of the DIA-MS Workflow.** (A) Retention time calibration obtained for a replicate of the ‘Matched Normal’ group. Pink dots correspond to peptides used for the calibration, and the black line to the non-linear calibration curve. (B) Rank plot showing the protein abundance of the 1,802 quantifiable protein groups. Abundances are depicted as median values. Pink dots correspond to matrisomal protein groups. (C-D) Boxplots of precursor abundance are displayed for each run (four runs associated with each patient) before (C) and after (D) local normalization.

**FIGURE S3: Proteomic Data Clustering - Assessment of Patient Gender.** Supervised clustering analysis using partial least squares-discriminant analysis (PLS-DA) performed on protein groups quantified in the ‘Matched Normal’ (shades of yellow) and ‘Tumor’ (shades of grey) samples collected on five female and five male patients with lung squamous cell carcinoma.

**FIGURE S4: Remodeling of the Matrisomal Proteins in Lung Squamous Cell Cancer.** List of the 66 protein groups significantly altered in ‘Tumor’ vs. ‘Matched Normal’, that are reported in the human MatrisomeDB [50].

**FIGURE S5: Heatmaps for Significantly Altered Serpins, Keratins and Desmosomal Proteins in LSCC across Individual Patients.** Heatmap showing log_2_-fold protein changes (‘Tumor’ vs. ’Matched Normal’) for each individual LSCC patient referred to as L01, L02, L03 …, and L10 assessing Core Protein Signatures. (A) SERPINB5 and (B) fifteen keratins were significantly up-regulated, while (A) SERPINB6 and (B) two keratins were significantly down-regulated in ‘Tumor’ vs. ‘Matched Normal’. In all displayed cases, Q-values (not displayed) were smaller than 2.24e-4, when comparing ‘Matched Normal’ group to ‘Tumor’ group (**Table S3**). (C) Schematic illustration of organization of keratins and desmosome proteins identified in the ECM fraction as insoluble remnants of stromal and epithelial cells.

## Notes

### Competing Interest Statement

The authors have declared no competing interest.

## References

[1] Tlsty, T. D., & Gascard, P. (2019). Stromal directives can control cancer. Science, 365(6449), 122–123.

[2] Zhao, H., Wu, L., Yan, G., Chen, Y., Zhou, M., Wu, Y., & Li, Y. (2021). Inflammation and tumor progression: signaling pathways and targeted intervention. Signal Transduct Target Ther, 6(1), 263.

[3] Qian, S., Golubnitschaja, O., & Zhan, X. (2019). Chronic inflammation: key player and biomarker-set to predict and prevent cancer development and progression based on individualized patient profiles. EPMA J, 10(4), 365–381.

[4] Hecht, S. S. (2011). Tobacco Smoke Carcinogens and Lung Cancer. In T. M. Penning (Ed.), Chemical Carcinogenesis (pp. 53–74). Totowa, NJ: Humana Press.

[5] Tsao, A. S., Liu, D., Lee, J. J., Spitz, M., & Hong, W. K. (2006). Smoking affects treatment outcome in patients with advanced nonsmall cell lung cancer. Cancer, 106(11), 2428–2436.

[6] Kenfield, S. A., Wei, E. K., Stampfer, M. J., Rosner, B. A., & Colditz, G. A. (2008). Comparison of aspects of smoking among the four histological types of lung cancer. Tob Control, 17(3), 198–204.

[7] Perez-Moreno, P., Brambilla, E., Thomas, R., & Soria, J. C. (2012). Squamous cell carcinoma of the lung: molecular subtypes and therapeutic opportunities. Clin Cancer Res, 18(9), 2443–2451.

[8] Jiang, Y., Zhang, H., Wang, J., Liu, Y., Luo, T., & Hua, H. (2022). Targeting extracellular matrix stiffness and mechanotransducers to improve cancer therapy. J Hematol Oncol, 15(1), 34.

[9] Henke, E., Nandigama, R., & Ergun, S. (2019). Extracellular Matrix in the Tumor Microenvironment and Its Impact on Cancer Therapy. Front Mol Biosci, 6, 160.

[10] Winkler, J., Abisoye-Ogunniyan, A., Metcalf, K. J., & Werb, Z. (2020). Concepts of extracellular matrix remodelling in tumour progression and metastasis. Nat Commun, 11(1), 5120.

[11] Naba, A., Clauser, K. R., Ding, H., Whittaker, C. A., Carr, S. A., & Hynes, R. O. (2016). The extracellular matrix: Tools and insights for the “omics” era. Matrix Biol, 49, 10–24.

[12] Naba, A., Clauser, K. R., Hoersch, S., Liu, H., Carr, S. A., & Hynes, R. O. (2012). The matrisome: in silico definition and in vivo characterization by proteomics of normal and tumor extracellular matrices. Mol Cell Proteomics, 11(4), M111 014647.

[13] McCabe, M. C., Saviola, A. J., & Hansen, K. C. (2022). A mass spectrometry-based atlas of extracellular matrix proteins across 25 mouse organs. Preprint. bioRxiv 2022.03.04.482898, https://doi.org/10.1101/2022.1103.1104.482898.

[14] Trombetta-Lima, M., Rosa-Fernandes, L., Angeli, C. B., Moretti, I. F., Franco, Y. M., Mousessian, A. S., . . . Palmisano, G. (2021). Extracellular Matrix Proteome Remodeling in Human Glioblastoma and Medulloblastoma. J Proteome Res, 20(10), 4693–4707.

[15] Moreira, A. M., Ferreira, R. M., Carneiro, P., Figueiredo, J., Osorio, H., Barbosa, J., . . . Seruca, R. (2022). Proteomic Identification of a Gastric Tumor ECM Signature Associated With Cancer Progression. Front Mol Biosci, 9, 818552.

[16] Tian, C., Clauser, K. R., Ohlund, D., Rickelt, S., Huang, Y., Gupta, M., . . . Hynes, R. O. (2019). Proteomic analyses of ECM during pancreatic ductal adenocarcinoma progression reveal different contributions by tumor and stromal cells. Proc Natl Acad Sci U S A, 116(39), 19609–19618.

[17] Hebert, J. D., Myers, S. A., Naba, A., Abbruzzese, G., Lamar, J. M., Carr, S. A., & Hynes, R. O. (2020). Proteomic Profiling of the ECM of Xenograft Breast Cancer Metastases in Different Organs Reveals Distinct Metastatic Niches. Cancer Res, 80(7), 1475–1485.

[18] Wishart, A. L., Conner, S. J., Guarin, J. R., Fatherree, J. P., Peng, Y., McGinn, R. A., . . . Oudin, M. J. (2020). Decellularized extracellular matrix scaffolds identify full-length collagen VI as a driver of breast cancer cell invasion in obesity and metastasis. Sci Adv, 6(43), eabc3175.

[19] Naba, A., Clauser, K. R., Mani, D. R., Carr, S. A., & Hynes, R. O. (2017). Quantitative proteomic profiling of the extracellular matrix of pancreatic islets during the angiogenic switch and insulinoma progression. Sci Rep, 7, 40495.

[20] Ross, P. L., Huang, Y. N., Marchese, J. N., Williamson, B., Parker, K., Hattan, S., . . . Pappin, D. J. (2004). Multiplexed protein quantitation in Saccharomyces cerevisiae using amine-reactive isobaric tagging reagents. Mol Cell Proteomics, 3(12), 1154–1169.

[21] Thompson, A., Schafer, J., Kuhn, K., Kienle, S., Schwarz, J., Schmidt, G., . . . Hamon, C. (2003). Tandem mass tags: a novel quantification strategy for comparative analysis of complex protein mixtures by MS/MS. Anal Chem, 75(8), 1895–1904.

[22] Ting, L., Rad, R., Gygi, S. P., & Haas, W. (2011). MS3 eliminates ratio distortion in isobaric multiplexed quantitative proteomics. Nat Methods, 8(11), 937–940.

[23] Wenger, C. D., Lee, M. V., Hebert, A. S., McAlister, G. C., Phanstiel, D. H., Westphall, M. S., & Coon, J. J. (2011). Gas-phase purification enables accurate, multiplexed proteome quantification with isobaric tagging. Nat Methods, 8(11), 933–935.

[24] Erickson, B. K., Rose, C. M., Braun, C. R., Erickson, A. R., Knott, J., McAlister, G. C., . . . Gygi, S. P. (2017). A Strategy to Combine Sample Multiplexing with Targeted Proteomics Assays for High-Throughput Protein Signature Characterization. Mol Cell, 65(2), 361–370.

[25] Gillet, L. C., Navarro, P., Tate, S., Rost, H., Selevsek, N., Reiter, L., . . . Aebersold, R. (2012). Targeted data extraction of the MS/MS spectra generated by data-independent acquisition: a new concept for consistent and accurate proteome analysis. Mol Cell Proteomics, 11(6), O111 016717.

[26] Ludwig, C., Gillet, L., Rosenberger, G., Amon, S., Collins, B. C., & Aebersold, R. (2018). Data-independent acquisition-based SWATH-MS for quantitative proteomics: a tutorial. Mol Syst Biol, 14(8), e8126.

[27] Bons, J., Rose, J., O’Broin, A., & Schilling, B. (2022). Advanced mass spectrometry-based methods for protein molecular-structural biologists. In Advances in Protein Molecular and Structural Biology Methods (pp. 311–326).

[28] Zhang, F., Ge, W., Ruan, G., Cai, X., & Guo, T. (2020). Data-Independent Acquisition Mass Spectrometry-Based Proteomics and Software Tools: A Glimpse in 2020. Proteomics, 20(17-18), e1900276.

[29] Bouchal, P., Schubert, O. T., Faktor, J., Capkova, L., Imrichova, H., Zoufalova, K., . . . Aebersold, R. (2019). Breast Cancer Classification Based on Proteotypes Obtained by SWATH Mass Spectrometry. Cell Rep, 28(3), 832–843 e7.

[30] Sun, R., Lyu, M., Liang, S., Ge, W., Wang, Y., Ding, X., . . . Guo, T. (2022). A prostate cancer tissue specific spectral library for targeted proteomic analysis. Proteomics, 22(7), e2100147.

[31] Rosenberger, G., Koh, C. C., Guo, T., Rost, H. L., Kouvonen, P., Collins, B. C., . . . Aebersold, R. (2014). A repository of assays to quantify 10,000 human proteins by SWATH-MS. Sci Data, 1, 140031.

[32] Krasny, L., Bland, P., Burns, J., Lima, N. C., Harrison, P. T., Pacini, L., . . . Huang, P. H. (2020). A mouse SWATH-mass spectrometry reference spectral library enables deconvolution of species-specific proteomic alterations in human tumour xenografts. Dis Model Mech, 13(7), dmm044586.

[33] Tsou, C. C., Avtonomov, D., Larsen, B., Tucholska, M., Choi, H., Gingras, A. C., & Nesvizhskii, A. I. (2015). DIA-Umpire: comprehensive computational framework for data-independent acquisition proteomics. Nat Methods, 12(3), 258–264, 257 p following 264.

[34] Demichev, V., Messner, C. B., Vernardis, S. I., Lilley, K. S., & Ralser, M. (2020). DIA-NN: neural networks and interference correction enable deep proteome coverage in high throughput. Nat Methods, 17(1), 41–44.

[35] Navarro, P., Kuharev, J., Gillet, L. C., Bernhardt, O. M., MacLean, B., Rost, H. L., . . . Tenzer, S. (2016). A multicenter study benchmarks software tools for label-free proteome quantification. Nat Biotechnol, 34(11), 1130–1136.

[36] Gotti, C., Roux-Dalvai, F., Joly-Beauparlant, C., Mangnier, L., Leclercq, M., & Droit, A. (2021). Extensive and Accurate Benchmarking of DIA Acquisition Methods and Software Tools Using a Complex Proteomic Standard. J Proteome Res, 20(10), 4801–4814.

[37] Collins, B. C., Hunter, C. L., Liu, Y., Schilling, B., Rosenberger, G., Bader, S. L., . . . Aebersold, R. (2017). Multi-laboratory assessment of reproducibility, qualitative and quantitative performance of SWATH-mass spectrometry. Nat Commun, 8(1), 291.

[38] Schilling, B., Gibson, B. W., & Hunter, C. L. (2017). Generation of High-Quality SWATH((R)) Acquisition Data for Label-free Quantitative Proteomics Studies Using TripleTOF((R)) Mass Spectrometers. Methods Mol Biol, 1550, 223–233.

[39] Meier, F., Brunner, A. D., Frank, M., Ha, A., Bludau, I., Voytik, E., . . . Mann, M. (2020). diaPASEF: parallel accumulation-serial fragmentation combined with data-independent acquisition. Nat Methods, 17(12), 1229–1236.

[40] Krasny, L., & Huang, P. H. (2021). Data-independent acquisition mass spectrometry (DIA-MS) for proteomic applications in oncology. Mol Omics, 17(1), 29–42.

[41] Meyer, J. G., & Schilling, B. (2017). Clinical applications of quantitative proteomics using targeted and untargeted data-independent acquisition techniques. Expert Rev Proteomics, 14(5), 419–429.

[42] Krasny, L., Bland, P., Kogata, N., Wai, P., Howard, B. A., Natrajan, R. C., & Huang, P. H. (2018). SWATH mass spectrometry as a tool for quantitative profiling of the matrisome. J Proteomics, 189, 11–22.

[43] Escher, C., Reiter, L., MacLean, B., Ossola, R., Herzog, F., Chilton, J., . . . Rinner, O. (2012). Using iRT, a normalized retention time for more targeted measurement of peptides. Proteomics, 12(8), 1111–1121.

[44] Burger, T. (2018). Gentle Introduction to the Statistical Foundations of False Discovery Rate in Quantitative Proteomics. J Proteome Res, 17(1), 12–22.

[45] Wickham, H. (2009). ggplot2: elegant graphics for data analysis: Springer New York.

[46] Rohart, F., Gautier, B., Singh, A., & Le Cao, K. A. (2017). mixOmics: An R package for ’omics feature selection and multiple data integration. PLoS Comput Biol, 13(11), e1005752.

[47] Kamburov, A., Pentchev, K., Galicka, H., Wierling, C., Lehrach, H., & Herwig, R. (2011). ConsensusPathDB: toward a more complete picture of cell biology. Nucleic Acids Res, 39(Database issue), D712–717.

[48] Kamburov, A., Wierling, C., Lehrach, H., & Herwig, R. (2009). ConsensusPathDB-- a database for integrating human functional interaction networks. Nucleic Acids Res, 37(Database issue), D623–628.

[49] Callister, S. J., Barry, R. C., Adkins, J. N., Johnson, E. T., Qian, W. J., Webb-Robertson, B. J., . . . Lipton, M. S. (2006). Normalization approaches for removing systematic biases associated with mass spectrometry and label-free proteomics. J Proteome Res, 5(2), 277–286.

[50] Shao, X., Taha, I. N., Clauser, K. R., Gao, Y. T., & Naba, A. (2020). MatrisomeDB: the ECM-protein knowledge database. Nucleic Acids Res, 48(D1), D1136–D1144.

[51] Midwood, K. S., Chiquet, M., Tucker, R. P., & Orend, G. (2016). Tenascin-C at a glance. J Cell Sci, 129(23), 4321–4327.

[52] Parekh, K., Ramachandran, S., Cooper, J., Bigner, D., Patterson, A., & Mohanakumar, T. (2005). Tenascin-C, over expressed in lung cancer down regulates effector functions of tumor infiltrating lymphocytes. Lung Cancer, 47(1), 17–29.

[53] Biaoxue, R., Xiguang, C., & Shuanying, Y. (2014). Annexin A1 in malignant tumors: current opinions and controversies. Int J Biol Markers, 29(1), e8–20.

[54] Elakad, O., Li, Y., Gieser, N., Yao, S., Kuffer, S., Hinterthaner, M., . . . Bohnenberger, H. (2021). Role of Annexin A1 in Squamous Cell Lung Cancer Progression. Dis Markers, 2021, 5520832.

[55] Oleksiewicz, U., Liloglou, T., Tasopoulou, K. M., Daskoulidou, N., Gosney, J. R., Field, J. K., & Xinarianos, G. (2017). COL1A1, PRPF40A, and UCP2 correlate with hypoxia markers in non-small cell lung cancer. J Cancer Res Clin Oncol, 143(7), 1133–1141.

[56] Dong, S., Zhu, P., & Zhang, S. (2020). Expression of collagen type 1 alpha 1 indicates lymph node metastasis and poor outcomes in squamous cell carcinomas of the lung. PeerJ, 8, e10089.

[57] Hou, L., Lin, T., Wang, Y., Liu, B., & Wang, M. (2021). Collagen type 1 alpha 1 chain is a novel predictive biomarker of poor progression-free survival and chemoresistance in metastatic lung cancer. J Cancer, 12(19), 5723–5731.

[58] Gonzalez-Gonzalez, L., & Alonso, J. (2018). Periostin: A Matricellular Protein With Multiple Functions in Cancer Development and Progression. Front Oncol, 8, 225.

[59] Kii, I., Nishiyama, T., Li, M., Matsumoto, K., Saito, M., Amizuka, N., & Kudo, A. (2010). Incorporation of tenascin-C into the extracellular matrix by periostin underlies an extracellular meshwork architecture. J Biol Chem, 285(3), 2028–2039.

[60] Ratajczak-Wielgomas, K., Kmiecik, A., Grzegrzolka, J., Piotrowska, A., Gomulkiewicz, A., Partynska, A., . . . Dziegiel, P. (2020). Prognostic Significance of Stromal Periostin Expression in Non-Small Cell Lung Cancer. Int J Mol Sci, 21(19).

[61] Ratajczak-Wielgomas, K., Kmiecik, A., & Dziegiel, P. (2022). Role of Periostin Expression in Non-Small Cell Lung Cancer: Periostin Silencing Inhibits the Migration and Invasion of Lung Cancer Cells via Regulation of MMP-2 Expression. Int J Mol Sci, 23(3).

[62] Warburg, O. (1956). On the origin of cancer cells. Science, 123(3191), 309–314.

[63] Liberti, M. V., & Locasale, J. W. (2016). The Warburg Effect: How Does it Benefit Cancer Cells? Trends Biochem Sci, 41(3), 211–218.

[64] Beloribi-Djefaflia, S., Vasseur, S., & Guillaumond, F. (2016). Lipid metabolic reprogramming in cancer cells. Oncogenesis, 5, e189.

[65] Vona, R., Iessi, E., & Matarrese, P. (2021). Role of Cholesterol and Lipid Rafts in Cancer Signaling: A Promising Therapeutic Opportunity? Front Cell Dev Biol, 9, 622908.

[66] Jayadev, R., & Sherwood, D. R. (2017). Basement membranes. Curr Biol, 27(6), R207–R211.

[67] Akashi, T., Ito, E., Eishi, Y., Koike, M., Nakamura, K., & Burgeson, R. E. (2001). Reduced expression of laminin alpha 3 and alpha 5 chains in non-small cell lung cancers. Jpn J Cancer Res, 92(3), 293–301.

[68] Chang, J., & Chaudhuri, O. (2019). Beyond proteases: Basement membrane mechanics and cancer invasion. J Cell Biol, 218(8), 2456–2469.

[69] Chen, S., & Birk, D. E. (2013). The regulatory roles of small leucine-rich proteoglycans in extracellular matrix assembly. FEBS J, 280(10), 2120–2137.

[70] Nastase, M. V., Iozzo, R. V., & Schaefer, L. (2014). Key roles for the small leucine-rich proteoglycans in renal and pulmonary pathophysiology. Biochim Biophys Acta, 1840(8), 2460–2470.

[71] Diehl, V., Huber, L. S., Trebicka, J., Wygrecka, M., Iozzo, R. V., & Schaefer, L. (2021). The Role of Decorin and Biglycan Signaling in Tumorigenesis. Front Oncol, 11, 801801.

[72] Appunni, S., Anand, V., Khandelwal, M., Gupta, N., Rubens, M., & Sharma, A. (2019). Small Leucine Rich Proteoglycans (decorin, biglycan and lumican) in cancer. Clin Chim Acta, 491, 1–7.

[73] Pietraszek-Gremplewicz, K., Karamanou, K., Niang, A., Dauchez, M., Belloy, N., Maquart, F. X., . . . Brezillon, S. (2019). Small leucine-rich proteoglycans and matrix metalloproteinase-14: Key partners? Matrix Biol, 75-76, 271–285.

[74] Kelly-Robinson, G. A., Reihill, J. A., Lundy, F. T., McGarvey, L. P., Lockhart, J. C., Litherland, G. J., . . . Martin, S. L. (2021). The Serpin Superfamily and Their Role in the Regulation and Dysfunction of Serine Protease Activity in COPD and Other Chronic Lung Diseases. Int J Mol Sci, 22(12).

[75] Zhao, X., Yuan, C., He, X., Wang, M., Zhang, H., Cheng, J., & Wang, H. (2022). Identification and in vitro validation of diagnostic and prognostic biomarkers for lung squamous cell carcinoma. J Thorac Dis, 14(4), 1243–1255.

[76] Ishida, Y., & Nagata, K. (2011). Hsp47 as a collagen-specific molecular chaperone. Methods Enzymol, 499, 167–182.

[77] Wang, Y., Gu, W., Wen, W., & Zhang, X. (2021). SERPINH1 is a Potential Prognostic Biomarker and Correlated With Immune Infiltration: A Pan-Cancer Analysis. Front Genet, 12, 756094.

[78] Zhu, J., Xiong, G., Fu, H., Evers, B. M., Zhou, B. P., & Xu, R. (2015). Chaperone Hsp47 Drives Malignant Growth and Invasion by Modulating an ECM Gene Network. Cancer Res, 75(8), 1580–1591.

[79] Xiong, G., Chen, J., Zhang, G., Wang, S., Kawasaki, K., Zhu, J., . . . Xu, R. (2020). Hsp47 promotes cancer metastasis by enhancing collagen-dependent cancer cell-platelet interaction. Proc Natl Acad Sci U S A, 117(7), 3748–3758.

[80] Nishimura, T., Fujii, K., Nakamura, H., Naruki, S., Sakai, H., Kimura, H., . . . Saji, H. (2021). Protein co-expression network-based profiles revealed from laser-microdissected cancerous cells of lung squamous-cell carcinomas. Sci Rep, 11(1), 20209.

[81] Park, H. J., Cha, Y. J., Kim, S. H., Kim, A., Kim, E. Y., & Chang, Y. S. (2017). Keratinization of Lung Squamous Cell Carcinoma Is Associated with Poor Clinical Outcome. Tuberc Respir Dis (Seoul), 80(2), 179–186.

[82] Galindo, I., Gomez-Morales, M., Diaz-Cano, I., Andrades, A., Caba-Molina, M., Miranda-Leon, M. T., . . . Farez-Vidal, M. E. (2020). The value of desmosomal plaque-related markers to distinguish squamous cell carcinoma and adenocarcinoma of the lung. Ups J Med Sci, 125(1), 19–29.

[83] Martin-Padron, J., Boyero, L., Rodriguez, M. I., Andrades, A., Diaz-Cano, I., Peinado, P., . . . Medina, P. P. (2020). Plakophilin 1 enhances MYC translation, promoting squamous cell lung cancer. Oncogene, 39(32), 5479–5493.

