## Supplementary figures and images for "Data-Independent Acquisition and Quantification of Extracellular Matrix from Human Lung in Chronic Inflammation-Associated Carcinomas"

### Figure S1

#### Supplementary Figure S1

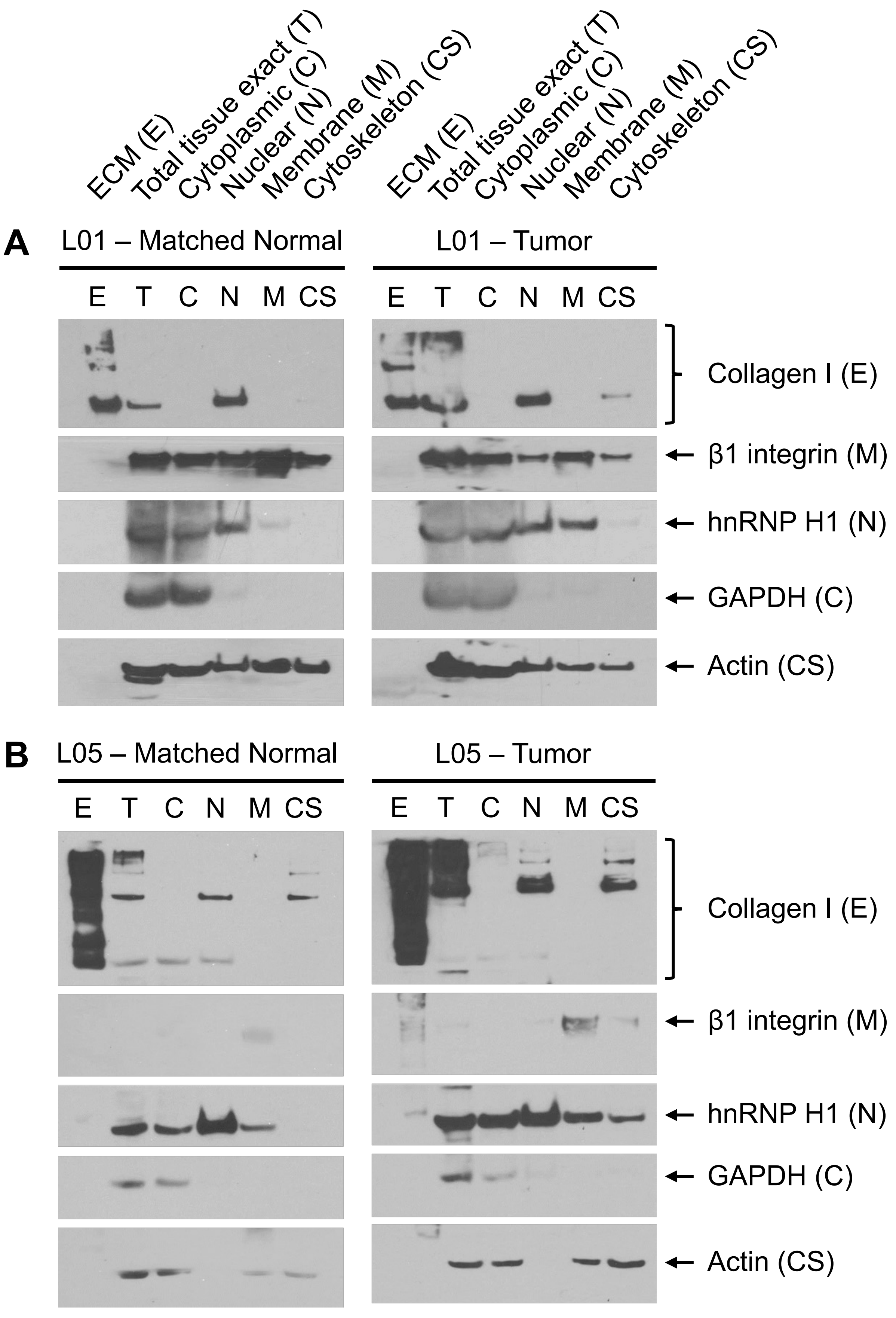

### Figure S2

#### Supplementary Figure S2

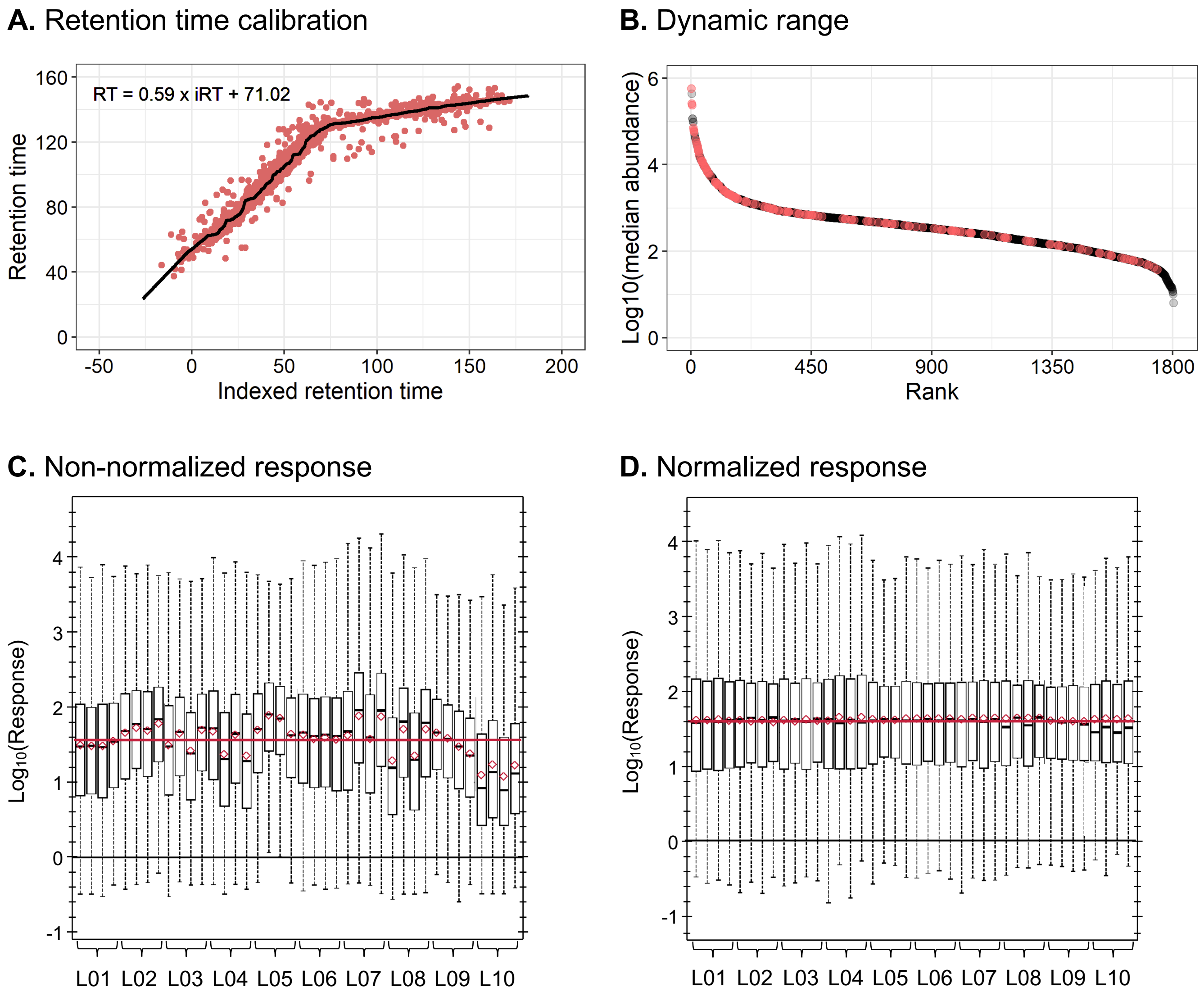

### Figure S3

#### Supplementary Figure S3

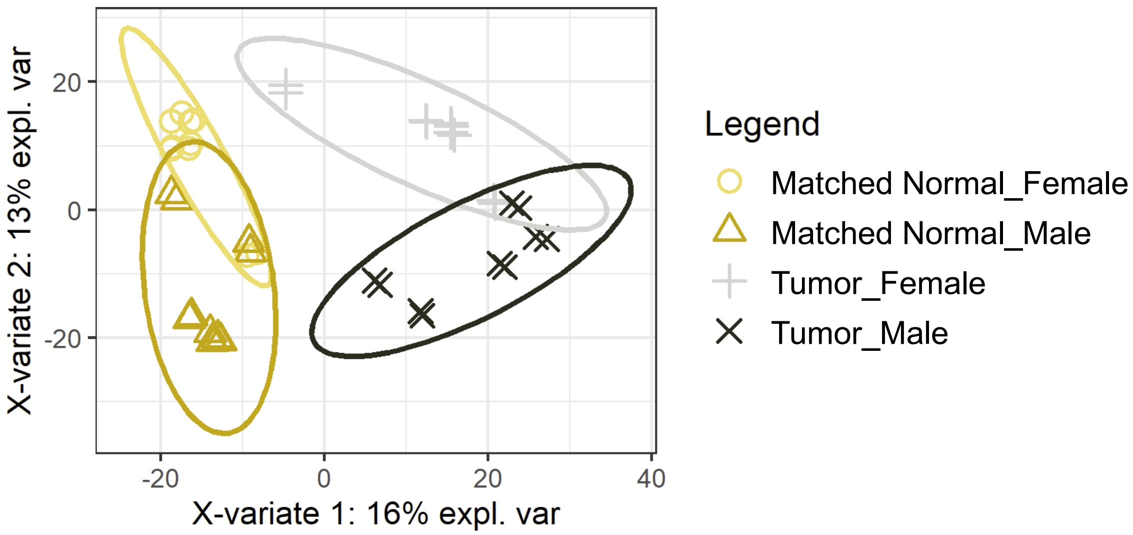
