## Supplementary material for "Data-Independent Acquisition and Quantification of Extracellular Matrix from Human Lung in Chronic Inflammation-Associated Carcinomas": Figure S4 - page 2

### Supplementary Figure S4

| Categories | Genes | Proteins | Ratio (Tumor vs Matched Normal) | $Log_2$ (Tumor vs Matched Normal) | Qvalue |
|---|---|---|---|---|---|
| Collagens | COL1A1 | Collagen alpha-1(I) chain | 3.25 | 1.70 | 8.81E-06 |
| | COL4A1 | Collagen alpha-1(IV) chain | 0.27 | -1.90 | 4.85E-09 |
| | COL4A2 | Collagen alpha-2(IV) chain | 0.21 | -2.22 | 1.44E-29 |
| | COL6A1 | Collagen alpha-1(VI) chain | 0.34 | -1.55 | 8.19E-77 |
| | COL6A2 | Collagen alpha-2(VI) chain | 0.34 | -1.55 | 8.22E-78 |
| | COL6A3 | Collagen alpha-3(VI) chain | 0.37 | -1.44 | < 1.00E-253 |
| | COL6A6 | Collagen alpha-6(VI) chain | 0.08 | -3.57 | 1.67E-15 |
| | COL7A1 | Collagen alpha-1(VII) chain | 4.01 | 2.00 | 2.12E-33 |
| | COL12A1 | Collagen alpha-1(XII) chain | 0.43 | -1.20 | 1.23E-23 |
| | COL14A1 | Collagen alpha-1(XIV) chain | 0.63 | -0.68 | 3.55E-80 |
| | COL15A1 | Collagen alpha-1(XV) chain | 0.63 | -0.68 | 3.24E-10 |
| | COL18A1 | Collagen alpha-1(XVIII) chain | 0.37 | -1.42 | 2.96E-36 |
| ECM glycoproteins | ABI3BP | Target of Nesh-SH3 | 0.47 | -1.10 | 5.17E-11 |
| | DPT | Dermatopontin | 0.47 | -1.10 | 7.76E-07 |
| | FBLN1 | Fibulin-1 | 0.60 | -0.73 | 1.18E-05 |
| | FBLN5 | Fibulin-5 | 0.42 | -1.26 | 3.74E-11 |
| | FGG | Fibrinogen gamma chain | 0.47 | -1.10 | 9.23E-04 |
| | LAMA3 | Laminin subunit alpha-3 | 0.29 | -1.77 | 6.36E-14 |
| | LAMA4 | Laminin subunit alpha-4 | 0.22 | -2.16 | 7.77E-76 |
| | LAMA5 | Laminin subunit alpha-5 | 0.18 | -2.49 | 6.28E-174 |
| | LAMB1 | Laminin subunit beta-1 | 0.41 | -1.30 | 5.07E-12 |
| | LAMB2 | Laminin subunit beta-2 | 0.16 | -2.64 | 5.61E-132 |
| | LAMB3 | Laminin subunit beta-3 | 0.17 | -2.56 | 1.18E-17 |
| | LAMC1 | Laminin subunit gamma-1 | 0.18 | -2.49 | 2.65E-112 |
| | LAMC2 | Laminin subunit gamma-2 | 0.24 | -2.09 | 1.47E-23 |
| | MATN2 | Matrilin-2 | 0.56 | -0.83 | 5.74E-07 |
| | MFAP4 | Microfibril-associated glycoprotein 4 | 0.40 | -1.32 | 1.81E-16 |
| | MMRN1 | Multimerin-1 | 0.44 | -1.20 | 7.73E-04 |
| | MMRN2 | Multimerin-2 | 0.35 | -1.52 | 6.00E-19 |
| | NID1 | Nidogen-1 | 0.14 | -2.80 | 5.56E-96 |
| | PAPLN | Papilin | 0.34 | -1.58 | 3.68E-05 |
| | POSTN | Periostin | 3.86 | 1.95 | 2.99E-64 |
| | SPON1 | Spondin-1 | 0.47 | -1.09 | 2.33E-09 |
| | TGFBI | Transforming growth factor-beta-induced protein ig-h3 | 0.66 | -0.61 | 2.96E-17 |
| | THBS1 | Thrombospondin-1 | 2.43 | 1.28 | 7.31E-07 |
| | TINAGL1 | Tubulointerstitial nephritis antigen-like | 0.36 | -1.48 | 5.38E-14 |
| | TNC | Tenascin | 4.16 | 2.06 | 2.57E-69 |
| | TNXB | Tenascin-X | 0.33 | -1.60 | 2.83E-55 |
| | VTN | Vitronectin | 0.42 | -1.25 | 3.21E-35 |
| | VWA1 | von Willebrand factor A domain-containing protein 1 | 0.34 | -1.55 | 3.32E-37 |
| | VWF | von Willebrand factor | 0.27 | -1.88 | 1.13E-42 |

| Categories | Genes | Proteins | Ratio (Tumor vs Matched Normal) | $Log_2$ (Tumor vs Matched Normal) | Qvalue |
|---|---|---|---|---|---|
| Proteoglycans | ASPN | Asporin | 0.55 | -0.87 | 4.42E-09 |
| | BGN | Biglycan | 0.30 | -1.76 | 5.35E-54 |
| | DCN | Decorin | 0.17 | -2.57 | 6.14E-30 |
| | HSPG2 | Basement membrane-specific heparan sulfate proteoglycan core protein | 0.17 | -2.56 | < 1.00E-253 |
| | LUM | Lumican | 0.18 | -2.48 | 2.77E-27 |
| | OGN | Mimecan | 0.09 | -3.44 | 4.90E-41 |
| | PRELP | Prolargin | 0.43 | -1.20 | 9.14E-27 |
| | PRG4 | Proteoglycan 4 | 0.43 | -1.21 | 9.38E-08 |
| ECM-affiliated proteins | ANXA1 | Annexin A1 | 4.06 | 2.02 | 5.27E-07 |
| | ANXA2 | Annexin A2 | 0.38 | -1.41 | 5.62E-42 |
| | ANXA6 | Annexin A6 | 0.62 | -0.69 | 6.03E-06 |
| | LGALS7 | Galectin-7 | 2.64 | 1.40 | 2.69E-04 |
| ECM regulators | AGT | Angiotensinogen | 0.54 | -0.89 | 5.93E-04 |
| | AMBP | Protein AMBP | 0.24 | -2.04 | 2.12E-26 |
| | CTSB | Cathepsin B | 2.07 | 1.05 | 1.09E-05 |
| | HRG | Histidine-rich glycoprotein | 0.53 | -0.92 | 7.04E-10 |
| | KNG1 | Kininogen-1 | 0.66 | -0.60 | 7.93E-04 |
| | MMP9 | Matrix metalloproteinase-9 | 0.66 | -0.61 | 8.77E-04 |
| | SERPINB5 | Serpin B5 | 1.91 | 0.94 | 8.20E-09 |
| | SERPINB6 | Serpin B6 | 0.59 | -0.75 | 2.24E-04 |
| | SERPINH1 | Serpin H1 | 2.35 | 1.23 | 7.84E-15 |
| | TGM2 | Protein-glutamine gamma-glutamyltransferase 2 | 0.48 | -1.07 | 5.63E-37 |
| Secreted factors | FGF2 | Fibroblast growth factor 2 | 0.39 | -1.35 | 9.07E-07 |
| | S100A11 | Protein S100-A11 | 2.40 | 1.26 | 1.32E-14 |
| | S100A9 | Protein S100-A9 | 1.91 | 0.93 | 1.49E-05 |
