## Supplementary material for "Data-Independent Acquisition and Quantification of Extracellular Matrix from Human Lung in Chronic Inflammation-Associated Carcinomas": Figure S5

### Supplementary Figure S5

**A.** Significantly altered serpin proteins

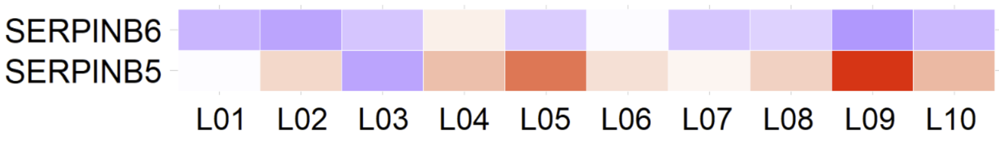


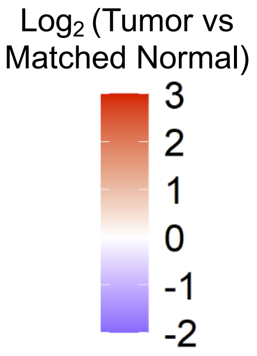


**B.** Significantly altered keratins

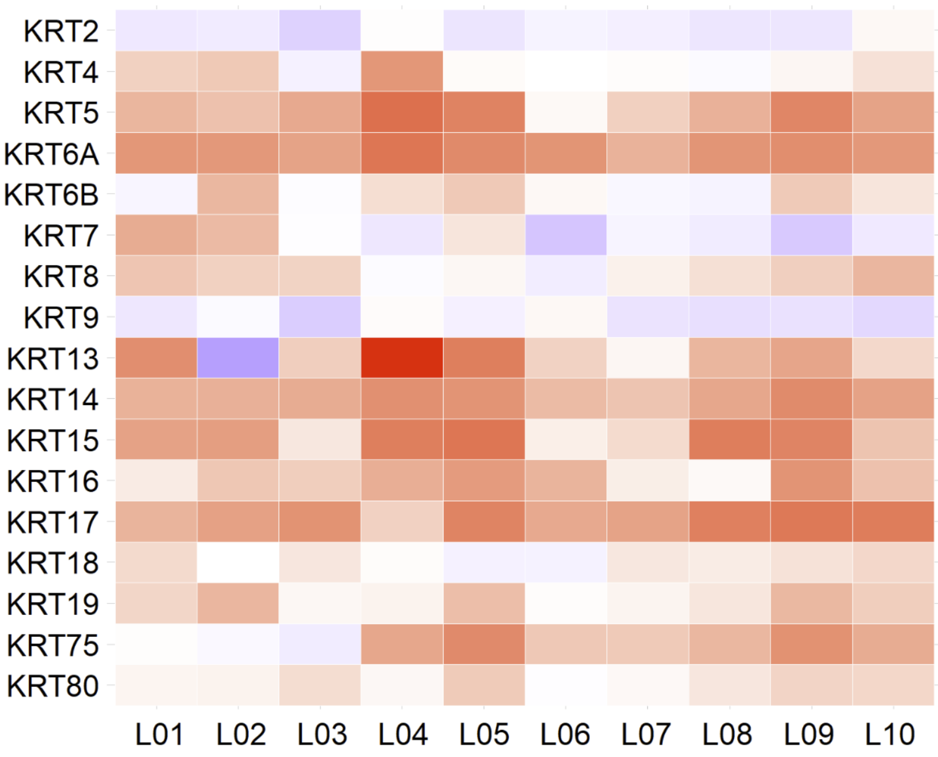


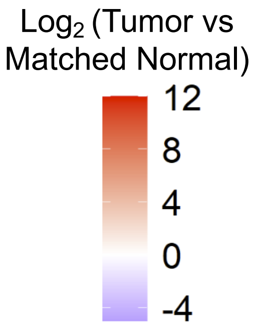


**C.** Schematic illustration of keratins and desmosome

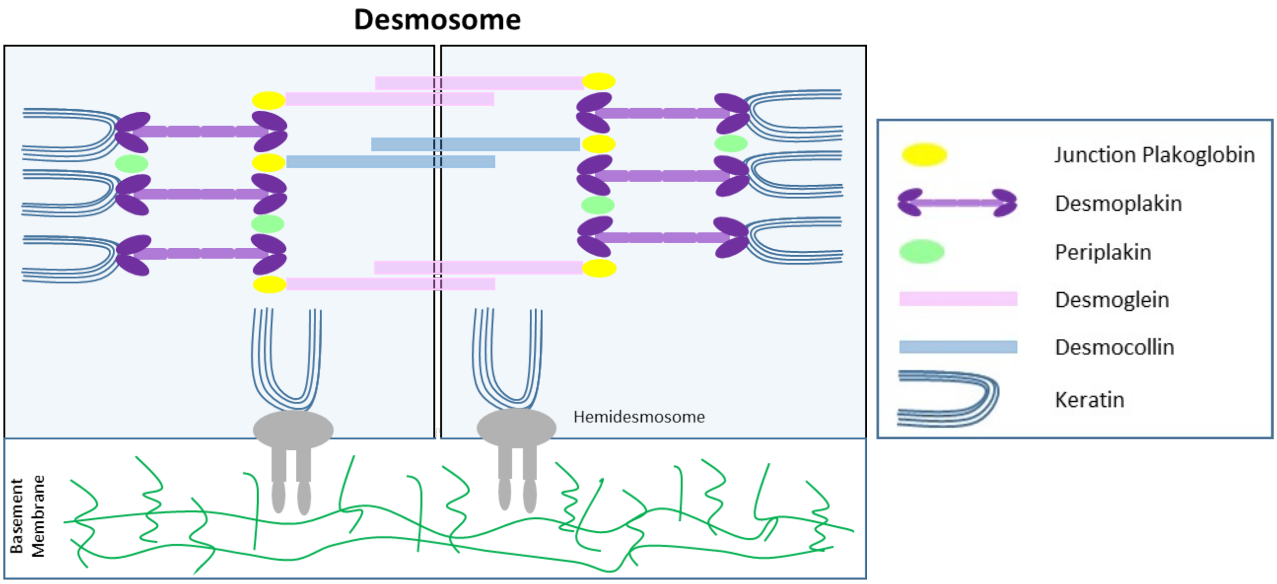
